# Beneficial effects of intermittent fasting in NASH and subsequent HCC development are executed by concerted PPARα and PCK1 action in hepatocytes

**DOI:** 10.1101/2023.10.23.562885

**Authors:** Suchira Gallage, Adnan Ali, Jose Efren Barragan Avila, Nogayhan Seymen, Pierluigi Ramadori, Vera Joerke, Laimdota Zizmare, Jan Kosla, Xin Li, Enrico Focaccia, Suhail Yousuf, Tjeerd Sijmonsma, Mohammad Rahbari, Katharina S. Kommoss, Adrian Billeter, Sandra Prokosch, Ulrike Rothermel, Florian Mueller, Jenny Hetzer, Danijela Heide, Tim Machauer, Nisar P. Malek, Thomas Longerich, Adam J. Rose, Susanne Roth, Johannes Schwenck, Christoph Trautwein, Mohammad M Karimi, Mathias Heikenwalder

## Abstract

The role and molecular mechanisms of intermittent fasting (IF) in non-alcoholic steatohepatitis (NASH) and its transition to hepatocellular carcinoma (HCC) are unknown. Here, we identified that an IF 5:2 regimen (two non-consecutive days of food deprivation per week), initiated in the active phase of mice, prevents/ameliorates NASH and fibrosis as well as reduces subsequent HCC development without affecting total calorie intake. The timing, length and number of fasting cycles as well as the type of NASH diet were all critical parameters determining the effectiveness of the fasting benefits. Combined proteomic, transcriptomic and metabolomic analyses identified that PPARα and glucocorticoid receptor (GR)-PCK1 act co-operatively as hepatic executors of the fasting response by promoting fatty acid catabolism and gluconeogenesis whilst suppressing anabolic lipogenesis. In line, PPARα targets and PCK1 were reduced in human NASH. Additionally, dynamic [^18^F]FDG-PET analysis *in vivo* revealed increased [^18^F]FDG uptake/retention and enhanced gluconeogenesis in the liver upon fasting (in accordance with PPARα and GR-PCK1 activation) when assessed by compartmental modelling. Hepatocyte-specific *GR* deletion only partially abrogated the hepatic fasting response. In contrast, the combined knockdown of *Ppara* and *Pck1 in vivo* abolished the beneficial outcomes of fasting against inflammation and fibrosis, confirming their causal relationship in integrating systemic signalling in hepatocytes. Notably, PPARα agonist pemafibrate recapitulated key aspects of hepatic fasting signalling at a molecular level. Therefore, IF or pharmacological mimetics of the PPARα and/or GR-PCK1 axis could be a viable intervention against NASH and subsequent liver cancer.

**One-Sentence Summary:** Intermittent fasting protects against fatty liver disease and liver cancer through concerted PPARα and GR-PCK1 action in hepatocytes.

## Main Text

Non-alcoholic fatty liver disease (NAFLD) is the most prevalent chronic liver disease worldwide and the incidence is expected to rise even further in the coming decades due to the obesity epidemic (Anstee et al., 2019). NAFLD is considered to be the hepatic manifestation of metabolic syndrome and can progress into more severe forms such as non-alcoholic steatohepatitis (NASH) and cirrhosis, associated with multiple complications and comorbidities (Anstee et al., 2019). Importantly, a subset of NASH patients can develop hepatocellular carcinoma (HCC), which is not only lethal but is the fastest-rising cancer in the USA and Europe with similar trends also observed in China and India (Gallage et al., 2021). The swift rise in the prevalence of obesity and associated pathologies including NASH in rapidly developing nations such as China can be partly attributed to a change in lifestyle from more traditional and balanced foods to more ultra-processed, high-caloric foods and sugary drinks, commonly termed the Western diet (Gallage et al., 2022b; Hall et al., 2019; Ma et al., 2021). Therefore, NASH/HCC imposes a huge present and future socioeconomic burden at a macroeconomic level but also diminishes the quality of life at an individual level. There are currently no approved pharmacological agents specifically targeting NAFLD, NASH or its transition to HCC (Ratziu et al., 2022). This raises the question of whether non-invasive, dietary restriction (DR)-based approaches can be utilized to treat this debilitating disease and if so, what cellular and molecular mechanisms would be the basis for possible beneficial effects.

In this light, intermittent fasting (IF) or other forms of dietary interventions such as time-restricted feeding (TRF) or fasting-mimicking diet (FMD) have gained increased popularity as viable interventions against obesity and metabolic diseases (Clifton et al., 2021; Longo et al., 2021). In addition, it was shown that mild caloric restriction in humans improved metabolic health and healthspan (Spadaro et al., 2022). Furthermore, IF and intermittent TRF (iTRF) have also been implicated to improve chronological healthspan and lifespan in flies (Catterson et al., 2018; Ulgherait et al., 2021). DR-based approaches have also been demonstrated to reduce body weight, blood pressure and atherogenic lipids in patients with metabolic syndrome as well as to modulate the gut microbiome to promote intestinal regeneration, reduce inflammatory bowel disease and alleviate diabetes-induced cognitive impairment (Liu et al., 2020; Rangan et al., 2019; Wilkinson et al., 2020). In some pre-clinical models, however, TRF was shown to not protect against atherosclerosis whilst IF showed beneficial effects against dyslipidemia and atherogenesis (Chen et al., 2021; Inoue et al., 2021; Mérian et al., 2023; Pan et al., 2023).

Given the increasing abundance of recent reports highlighting the benefits of fasting-based approaches, it is not surprising that this has trickled down to the general public as well. For example, the IF (5:2) regimen, whereby individuals voluntarily abstain (or significantly limit their calories) from intake of foods and calorie-dense liquids for two non-consecutive days a week, was the most popular dietary intervention in 2020 (Clifton et al., 2021). Although there is mounting evidence highlighting the benefits of DR-based interventions against obesity and non-liver-related pathologies, a potential benefit of fasting approaches in NASH and subsequent HCC development as well as the cellular and molecular mechanisms remain unknown. Therefore, we investigated the therapeutic potential of IF regimens in NASH and NASH to HCC transition using distinct obesogenic diet-induced models of NASH.

### Intermittent fasting protects against diet-induced obesity and NASH development

To investigate whether IF can prevent NASH development, as prompted by recent preliminary pre-clinical evidence that fasting can affect liver lipid metabolism to promote metabolic health (Chaix et al., 2021; Fuhrmeister et al., 2016), we utilized a well-characterized Western diet (WD)-based preclinical model of NASH that is rich in non-*trans* fats and sugars (fructose and sucrose) (Gallage et al., 2022a; Gallage et al., 2022b). 8-week-old male C57BL/6J mice were fed either normal chow diet (ND) or WD for 32 weeks to induce NASH (Figure 1A). The control groups (ND or WD) were given *ad libitum* (A/L) access to food and water (Figure 1A). In contrast, a group of mice fed WD underwent an IF (5:2) regimen involving two non-consecutive days of fasting (food deprivation but free access to water) per week with each fasting cycle lasting 24 hours (Figure 1A). Since mice are primarily nocturnal, the fasting cycle was initiated at the start of the active phase (7pm). Mice undergoing the IF (5:2) regimen had free access to food on all non-fasting days (Figure 1A). Of note, all of the physiological measurements (*e.g.,* body weight) as well as end-point measurements (*e.g.,* serology and histology) were made 48 hours after the last fasting cycle, which ensured all groups were at the same nutritional/feeding state (fed status) when analyzed unless stated otherwise.

**Figure 1.**
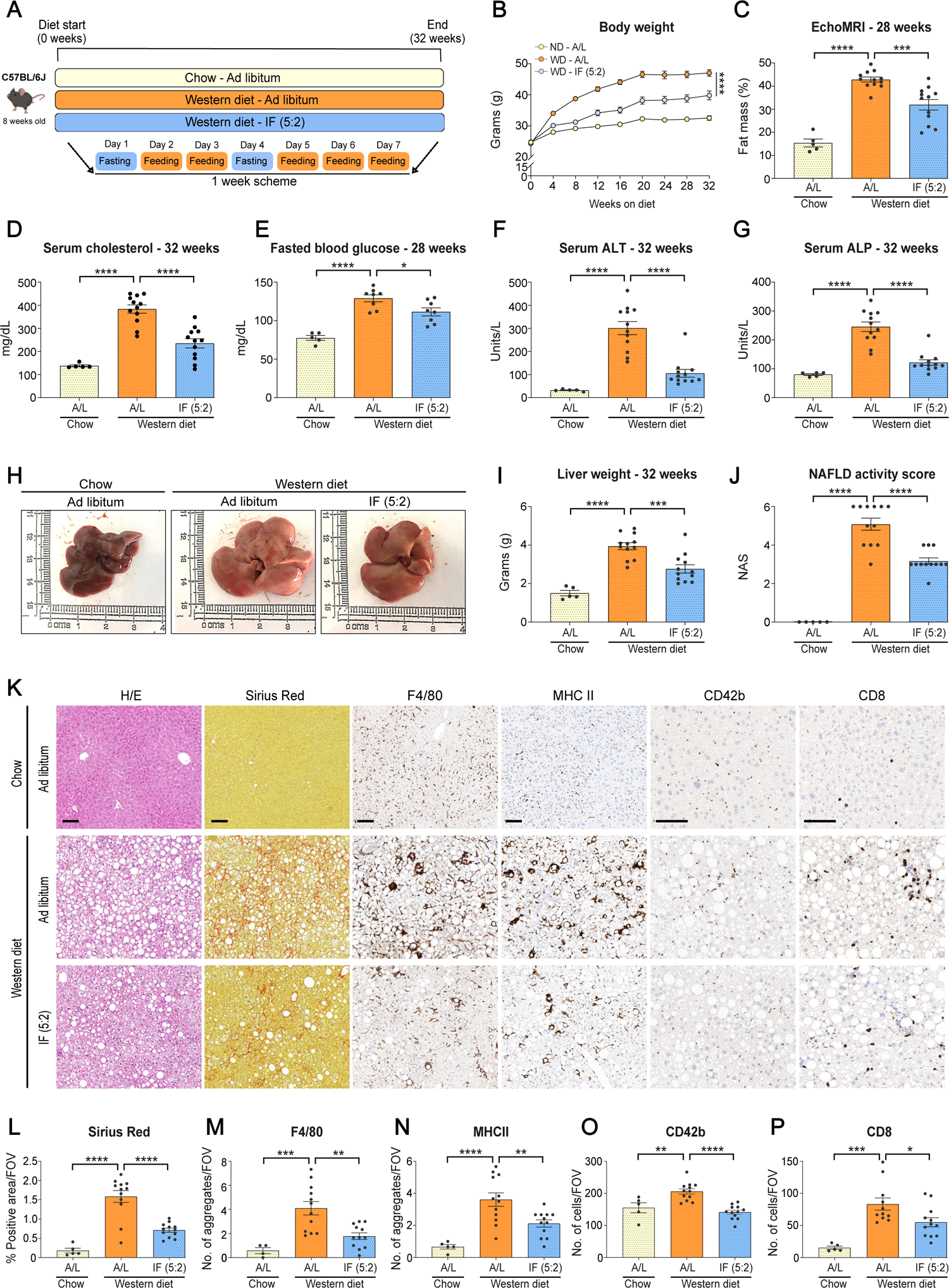
Intermittent fasting attenuates diet-induced obesity and NASH upon Western diet feeding in mice. **A.** Experimental scheme and graphical representation of the weekly fasting regimen. 8-week-old male C57BL/6J mice were fed with either normal chow diet or a Western diet (WD) for 32 weeks to induce NASH. The control groups (Chow or WD) were given *ad libitum* (A/L) access to food and water. In contrast, a group of mice fed WD underwent an IF (5:2) regimen involving two non-consecutive days of fasting per week with each fasting cycle lasting 24 hours. **B.** Bodyweight development over a period of 32 weeks. Chow A/L (n=5), WD A/L (n=12) and WD IF (5:2) (n=12). **C**. Fat mass determined by echoMRI. Chow A/L (n=5), WD A/L (n=12) and WD IF (5:2) (n=12). **D.** Serum levels of cholesterol at 32 weeks. Chow A/L (n=5), WD A/L (n=12) and WD IF (5:2) (n=12). **E**. 16h fasting serum glucose levels at 28 weeks post-start of experiment. Chow A/L (n=5), WD A/L (n=8) and WD IF (5:2) (n=8). **F and G.** Serum levels of alanine aminotransferase transaminase (ALT) and alkaline phosphatase (ALP) at 32 weeks. Chow A/L (n=5), WD A/L (n=12) and WD IF (5:2) (n=12). **H.** Representative macroscopic liver images from mice**. I.** Absolute liver weight in grams. Chow A/L (n=5), WD A/L (n=12) and WD IF (5:2) (n=12). **J.** NAS – NAFLD activity score assessing hepatic steatosis, ballooning and inflammation. Chow A/L (n=5), WD A/L (n=12) and WD IF (5:2) (n=12). **K.** Representative immunohistochemistry images for H/E Sirius Red, F4/80, MHCII, CD42b and CD8 staining from C57BL/6J mice fed a ND or WD under A/L conditions or WD with IF regimen for 32-weeks. **L-P.** Quantification of the indicated stainings. Sirius Red, MHCII and CD42b. Chow A/L (n=5), WD A/L (n=12) and WD IF (5:2) (n=12). F4/80 - Chow A/L (n=4), WD A/L (n=12) and WD IF (5:2) (n=12). CD8 – Chow A/L (n=5), WD A/L (n=11) and WD IF (5:2) (n=12). Data are expressed as mean ± SEM. Statistical significance was calculated using one-way analysis of variance with Tukey’s multiple comparison test. * P < 0.05, ** P < 0.01, *** P < 0.001, **** P < 0.0001. N.S: non-significant. Scale bar, 100µm.

WD-fed mice with A/L access to food gained significantly higher body weight with increased fat mass measured by EchoMRI as well as increased epididymal and inguinal white adipose tissue (eWAT/iWAT) depots compared to ND-fed mice (Figures 1B, 1C, and S1A-S1F). Whereas, mice undergoing the IF (5:2) regimen were resistant to WD-induced obesity with significantly lower body weight and body fat mass compared to WD-fed A/L controls (Figures 1B, 1C, and S1A-S1F). Importantly, fasted mice showed a significantly higher lean mass/body weight (%) compared to WD-fed A/L controls, indicating that the IF (5:2) regimen does not cause muscle wasting (Figures S1C and S1D). In addition, fasted mice displayed significantly lower levels of serum cholesterol and fasting blood glucose compared with WD-fed A/L controls, suggesting improved lipid and glucose homeostasis in the fasted mice (Figures 1D and 1E). Mice undergoing the IF (5:2) regimen not only showed metabolic improvements but also ameliorated liver pathology with strikingly lower serum levels of liver damage markers (ALT and ALP) as well as significantly lower liver weight (g) and liver to body weight ratio (%), indicating attenuated WD-induced hepatomegaly (Figures 1F-1I and S1G). Moreover, fasted mice showed significant improvements in hepatic steatosis, degenerative ballooning and inflammation as depicted by a significantly lower NAFLD activity score (NAS) compared to WD-fed A/L controls (Figure 1J and 1K). Finally, fasted mice also displayed significantly lower liver fibrosis and inflammation, specifically of myeloid aggregates (F4/80^+^ cells, MHCII^+^ cells), platelets (CD42b^+^ cells) and infiltration of auto-aggressive cytotoxic CD8^+^/PD1^+^ T-cells compared with WD-fed A/L controls (Figures 1K-1P and S1H). Overall, these findings indicate that the IF (5:2) regimen not only attenuates diet-induced obesity and metabolic dysfunction but NASH development as well.

### Fasting-mediated benefits are independent of total calorie intake

Upon observing that the IF (5:2) regimen attenuates NASH development, we then investigated whether this regimen affects total calorie intake and/or whole-body metabolism in mice fed a WD (Figure 2A). Interestingly, mice undergoing the IF (5:2) regimen did not show any significant differences in the average daily food intake (kcal/mouse/day) over the course of a week compared to either ND-fed or WD-fed A/L controls (Figure 2B). This was because the fasted mice displayed a hyperphagic response with significantly higher food intake in the feeding days, which compensated for the lack of food in the fasting days (Figures 2C, 2D, and S2A). This is consistent with previous reports showing that certain dietary regimens such as TRF also do not affect total calorie intake but still improve obesity and metabolic dysfunction (Chaix et al., 2019; Chaix et al., 2014). Additionally, there were no significant differences in the average daily water intake between

**Figure 2.**
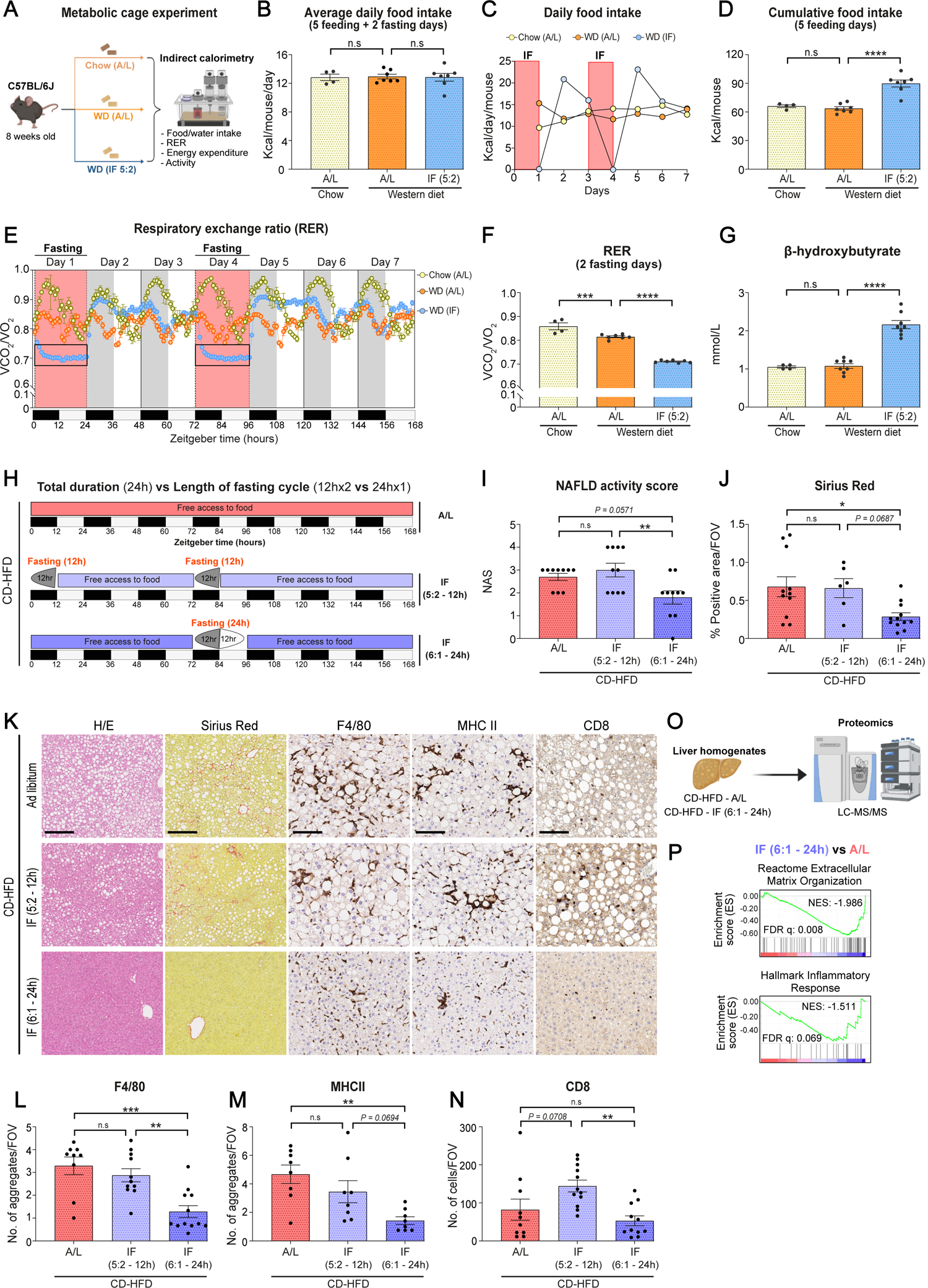
Fasting-mediated benefits are independent of total calorie intake but dependent on the length of the fasting cycle. **A.** Experimental scheme. Indirect calorimetry was carried out in 8-week-old male C57BL/6J mice to assess whole body metabolism. Mice were fed either a normal chow diet or a Western diet (WD). The control groups (Chow or WD) were given *ad libitum* (A/L) access to food and water. Mice under IF (5:2) regimen were fasted for two non-consecutive days per week with each fasting cycle lasting 24 hours. **B.** Average daily food intake over the span of 7 days in chow-fed A/L mice (n=4), WD-fed A/L mice (n=7) and WD-fed mice undergoing IF regimen (n=7). **C.** Daily food intake assessed over the span of 7 days. Chow A/L (n=4), WD A/L (n=7) and WD IF (5:2) (n=7). **D.** Cumulative food intake during the 5 feeding days. Chow A/L (n=4), WD A/L (n=7) and WD IF (5:2) (n=7). **E.** Respiratory exchange ratio (RER) over the span of 7 days in chow-fed A/L mice (n=4), WD-fed A/L mice (n=7) and WD-fed mice undergoing IF regimen (n=7). **F.** RER during the 2 fasting days. Chow A/L (n=4), WD A/L (n=7) and WD IF (5:2) (n=7). **G.** Serum levels of ketone body β-hydroxybutyrate in either chow (n=4) or WD (n=8) A/L mice under a fed status or in WD-fed mice undergoing IF 5:2 regimen (n=8) following a 24-hour fast. **H.** Experimental scheme. 8-week-old male C57BL/6J mice were fed a choline-deficient high-fat diet (CD-HFD) for 32 weeks to induce NASH. CD-HFD A/L mice was given free access to food and water. Mice under CD-HFD (5:2 - 12h) regimen were fasted for two non-consecutive days per week with each fasting cycle lasting 12 hours from 7pm to 7am. Mice under IF (6:1 - 24h) regimen were fasted for one day per week from 7pm to 7pm with each fasting cycle lasting 24 hours. **I**. NAS – NAFLD activity score assessing hepatic steatosis, ballooning and inflammation. CD-HFD A/L (n=10), CD-HFD IF (5:2 - 12h) (n=10) and CD-HFD IF (6:1 - 24h) (n=10). **J.** Quantification of Sirius red staining. CD-HFD A/L (n=11), CD-HFD IF (5:2 - 12h) (n=6) and CD-HFD IF (6:1 - 24h) (n=12). **K.** Representative immunohistochemistry images for H/E Sirius Red, F4/80, MHCII and CD8 staining from the indicated groups fed a CD-HFD. **L-N.** Quantification of corresponding stainings. F4/80 - CD-HFD A/L (n=9), CD-HFD IF (5:2 - 12h) (n=11) and CD-HFD IF (6:1 - 24h) (n=12). MHCII - CD-HFD A/L (n=8), CD-HFD IF (5:2 - 12h) (n=8) and CD-HFD IF (6:1 - 24h) (n=8). CD8 - CD-HFD A/L (n=10), CD-HFD IF (5:2 - 12h) (n=11) and CD-HFD IF (6:1 - 24h) (n=11). **O.** Graphical scheme. Proteomics analysis of the liver homogenates from CD-HFD A/L or CD-HFD IF (6:1) mice were carried out. **P.** Gene Set Enrichment Analysis (GSEA) for ‘REACTOME Extracellular’ and ‘HALLMARK Inflammatory Response’ in fasted mice compared to A/L fed controls. Data are expressed as mean ± SEM. Statistical significance was calculated using one-way analysis of variance with Tukey’s multiple comparison test. * P < 0.05, ** P < 0.01, *** P < 0.001, **** P < 0.0001. N.S: non-significant. Scale bar, 100µm.

WD-fed A/L or fasted mice (Figure S2B). We then assessed the impact of fasting on whole-body metabolism and observed that the respiratory exchange ratio (RER) was significantly lower (∼0.7) in fasted mice during the 2 fasting days compared to WD-fed A/L controls, indicating increased fatty acid oxidation (Figures 2E, 2F, S2C and S2D). Importantly, β-hydroxybutyrate (ketone body) a by-product of fatty acid oxidation and ketogenesis (Newman and Verdin, 2014), was significantly increased following a 24-hour fast (Figure 2G). Mice undergoing the IF (5:2) regimen also exhibited significantly lower oxygen consumption (VO_2_) and activity in the 2 fasting days compared to WD-fed A/L controls, while showing a tendency towards increased VO_2_ and activity in the feeding days (Figures S2E-S2J). Overall, mice undergoing the IF (5:2) regimen did not show any significant differences in the total calorie intake but displayed altered whole-body metabolism with an increased propensity towards fatty acid oxidation and ketogenesis during the fasting cycles despite decreased VO_2_ and activity.

### The length, number and timing of fasting cycles as well as the type of NASH diet determine the effectiveness of fasting benefits

We then systematically investigated which parameters are critical for an effective fasting regimen to yield benefits against NASH pathogenesis. We first aimed to understand whether the total duration of fasting per week or the length of the fasting cycle is more important in mediating the beneficial outcomes of fasting. To tackle this, we fed 8-week-old male C57BL/6J mice with the well-characterized choline-deficient high-fat diet (CD-HFD) model of NASH for 32 weeks (Figure 2H). We initially utilized the CD-HFD model since it is a milder NASH diet that presents with only mild fibrosis compared to the WD (Wolf et al., 2014). This enabled us to dissect the intricacies of the fasting regimens better. The control group (CD-HFD A/L) was given free access to food and water (Figure 2H). In contrast, mice were divided into two distinct IF regimens and were fed CD-HFD for 32 weeks as well (Figure 2H). In the first group termed the IF (5:2 - 12h) regimen, mice fed CD-HFD were fasted for two non-consecutive days of fasting per week with each fasting cycle lasting only 12 hours (Figure 2H). Importantly, each fasting cycle was initiated at the start of the active phase (7pm) and ended the following morning (7am). Whereas in the second group termed the IF (6:1 - 24h) regimen, mice fed CD-HFD were fasted for only one day per week but with each fasting cycle lasting 24 hours (Figure 2H). The fasting cycle was initiated at the start of the active phase (7pm) and ended the following evening (7pm). Therefore, both the IF groups were fasted for a total duration of 24 hours per week but with the IF (5:2 - 12h) group fasting twice per week for 12 hours while the IF (6:1 - 24h) group fasting only once per week for 24 consecutive hours.

Interestingly, mice undergoing the IF (6:1 - 24h) but not the IF (5:2 - 12h) displayed significantly lower body weight, body fat mass as well as lower levels of serum cholesterol and fasted blood glucose compared to CD-HFD-fed A/L controls (Figures S2K-S2P). However, both the IF (6:1 - 24h) and IF (5:2 - 12h) groups showed lower liver weight (g), liver/body weight (%) as well as lower serum ALT and ALP levels compared to CD-HFD-fed A/L controls, but with a more pronounced effect in the IF (6:1 - 24h) group (Figures S2Q-S2T). In line with this, mice undergoing the IF (6:1 - 24h) regimen showed strikingly lower hepatic steatosis, fibrosis and immune infiltration (myeloid: F4/80^+^ and MHCII^+^) compared to CD-HFD-fed A/L controls, which summated to a lower NAS (Figures 2I an 2N). Proteomics analyses corroborated the reduction in collagen deposition/fibrosis and immune infiltration in mice undergoing the IF (6:1 - 24h) regimen compared to A/L controls (Figures 2O and 2P). Mice undergoing the IF (5:2 - 12h) regimen, however, did not show significantly lower fibrosis, inflammation or NAS compared to CD-HFD-fed A/L controls (Figures 2K-2N). As before, neither the IF (6:1 - 24h) nor the IF (5:2 - 12h) regimen significantly affected total calorie intake but displayed reduced RER during the fasting period, indicative of increased fatty acid oxidation, compared to A/L controls (Figures S3 and S4). Therefore, these data demonstrate that the IF (6:1 - 24h) regimen induces a more beneficial outcome not only in the context of systemic obesity and metabolic parameters but also in liver pathology compared to the IF (5:2 - 12h) regimen in the context of CD-HFD feeding. This indicates that the length of the fasting cycle (24 consecutive hours > 12 hours performed twice) plays a more prominent role when keeping the total duration of fasting per week (24 hours) the same between groups.

To delve into the intricacies of the fasting cycle in more detail, we aimed to understand whether the number of fasting cycles per week as well as the timing of fasting (active phase vs inactive phase) also play a prominent role in mediating the benefits in NASH development. 8-week-old male C57BL/6J mice were fed with WD for 32 weeks to induce NASH (Figure 3A). The control groups (ND or WD) were given A/L access to food and water (Figure 3A). One group of mice was subjected to an IF (6:1) regimen consisting of fasting only once per week in the active phase (7pm to 7pm – 24 consecutive hours) (Figure 3A), akin to what was carried out in the context of a CD-HFD as well. With this, we aimed to determine whether one fasting cycle per week is sufficient to provide benefits in the context of a WD-induced NASH model as well. Importantly, WD is a more aggressive diet compared to CD-HFD, which only presents with mild fibrosis (Wolf et al., 2014). Therefore, depending on the severity of the diet, more fasting cycles may be needed to attain the benefits of fasting. In addition, mice fed with WD were also subjected to two additional fasting regimens: IF (6:1) or IF (5:2) initiated in the inactive phase (7am to 7am – 24 consecutive hours) (Figure 3A). Since mice are primarily nocturnal, this would be akin to initiating the fasting cycle in the evening in humans when most people would be sleeping. This is in contrast to the original experiment (Figure 1) where mice underwent the IF (5:2) regimen performed in the active phase (7pm to 7pm – 24 consecutive hours). This would allow us to determine if the timing of fasting initiation also affects the benefits.

**Figure 3.**
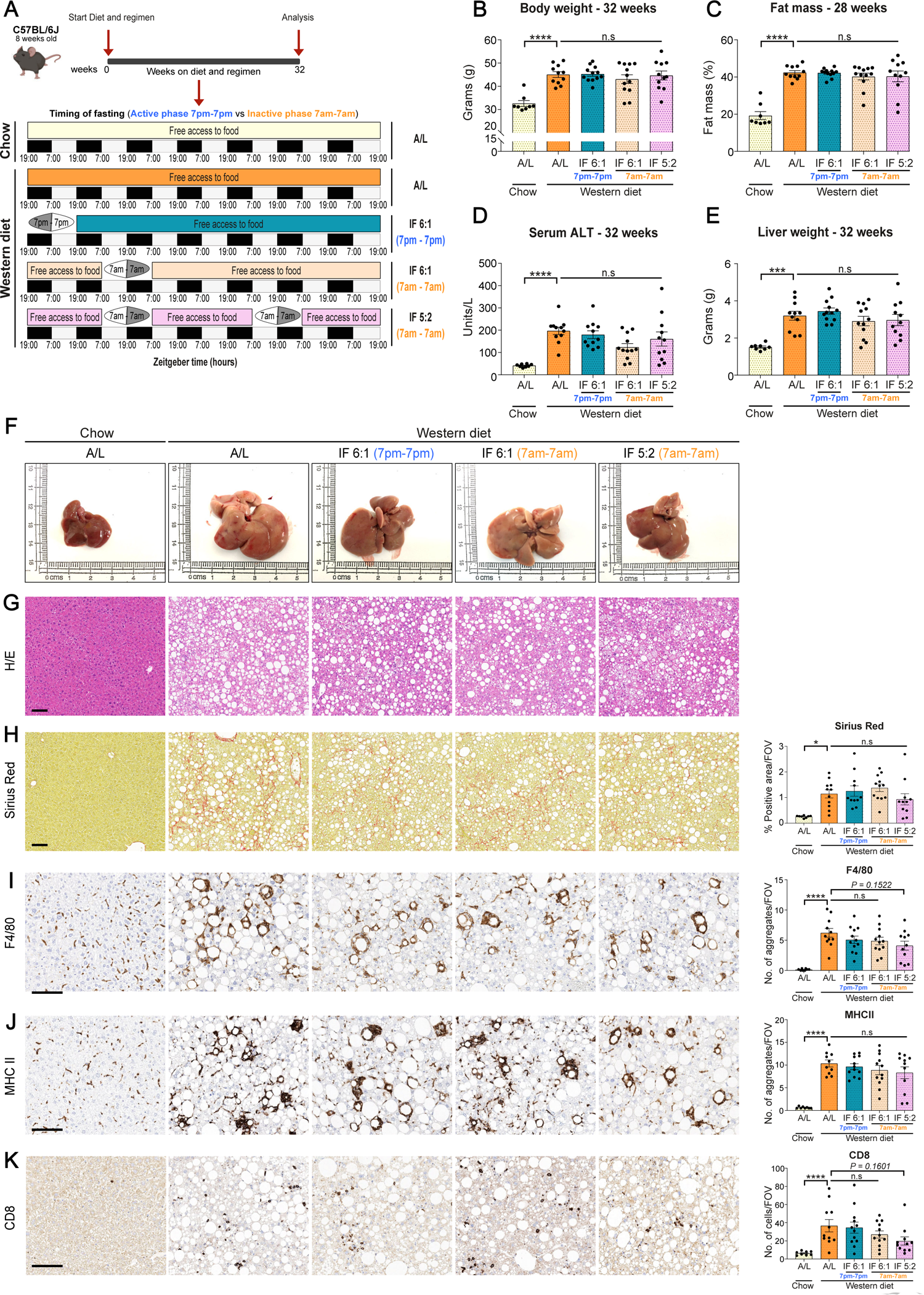
The timing and number of fasting cycles dictate the benefits of fasting in NASH pathogenesis. **A.** Experimental scheme. 8-week-old male C57BL/6J mice were fed a Western diet (WD) for 32 weeks to induce NASH. Chow and WD A/L mice was given free access to food and water. Mice under WD were also subjected to distinct fasting regimens. Mice under IF (6:1 - 7pm – 7pm) regimen were fasted for one day per week from 7pm to 7pm with each fasting cycle lasting 24 hours. Mice under IF (6:1 - 7am – 7am) regimen were fasted for one day per week from 7am to 7am with each fasting cycle lasting 24 hours. Mice under (5:2 – 7am – 7am) regimen were fasted for two non-consecutive days per week from 7am to 7am with each fasting cycle lasting 24 hours. **B.** Bodyweight at 32 weeks. Chow A/L (n=8), WD A/L (n=12), WD IF (6:1 - 7pm – 7pm) (n=11), WD IF (6:1 - 7am – 7am) (n=12) and WD IF (5:2 – 7am – 7am) (n=11). **C.** Fat mass determined by echoMRI at 28 weeks. Chow A/L (n=8), WD A/L (n=12), WD IF (6:1 - 7pm – 7pm) (n=11), WD IF (6:1 - 7am – 7am) (n=12) and WD IF (5:2 – 7am – 7am) (n=11). **D.** Serum levels of alanine aminotransferase transaminase (ALT) at 32 weeks. Chow A/L (n=8), WD A/L (n=11), WD IF (6:1 - 7pm – 7pm) (n=11), WD IF (6:1 - 7am – 7am) (n=12) and WD IF (5:2 – 7am – 7am) (n=11). **E.** Absolute liver weight in grams. Chow A/L (n=8), WD A/L (n=11), WD IF (6:1 - 7pm – 7pm) (n=12), WD IF (6:1 - 7am – 7am) (n=12) and WD IF (5:2 – 7am – 7am) (n=11). **F. Representative** macroscopic liver images of the indicated mice**. G.** Representative immunohistochemistry images for H/E of C57BL/6J mice fed Chow or WD under A/L conditions or WD with IF regimens for 32-weeks. **H-K.** Representative immunohistochemistry images (left) and quantification (right) for Sirius Red, F4/80, MHCII and CD8 staining of the indicated groups. Data are expressed as mean ± SEM. Statistical significance was calculated using one-way analysis of variance with Tukey’s multiple comparison test. * P < 0.05, *** P < 0.001, **** P < 0.0001. N.S: non-significant. Scale bar, 100µm.

Surprisingly, none of these fasting regimens (IF 6:1 active phase or IF 6:1/IF 5:2 inactive phase) displayed significant alterations in body weight, fat mass, lean mass nor in fasted blood glucose levels compared to WD-fed A/L controls (Figures 3B, 3C, and S5A-S5F). In addition, there were also no significant differences in serum liver damage markers (ALT and ALP) or in either the liver weight (g) or liver to body weight ratio (%) in mice undergoing the fasting regimens (IF 6:1 active phase or IF 6:1/IF 5:2 inactive phase) compared to WD-fed A/L controls (Figures 3D-3F, S5G, and S5H). These data indicate that these specific fasting regimens do not attenuate WD-induced metabolic dysfunction or hepatomegaly. Interestingly, the IF (5:2) regimen performed in the inactive phase showed a trend towards less (albeit not statistically significant) fibrosis and inflammation compared to WD-fed A/L controls, but none of the IF (6:1) regimens performed either in the active or inactive phase showed any significant improvements (Figures 3G-3K). Importantly, the IF (5:2) regimen performed in the inactive phase (7am to 7am) did not display the same striking diminution in either the metabolic parameters or liver pathology compared to when the IF (5:2) regimen was performed during the active phase (7pm to 7pm) (Figure S5I).

In summary, these data imply that there are four critical parameters affecting the efficacy of fasting in NASH (Figure S5I). Not only do timing (active phase > inactive phase) and length of the fasting cycle (24h > 12h*2) play an important role, but also the number of fasting cycles (2 cycles > 1 cycle per week) and the diet composition (the more aggressive WD needing more fasting cycles than the milder CD-HFD) in mediating the benefits of fasting in NASH (Figure S5I).

### Unbiased proteomics reveals PPARα and fatty acid oxidation at the center of the fasting **response during NASH.**

To determine the mechanistic basis underlying the benefits of IF in NASH, we fed C57BL/6J mice with ND (chow) or WD for 5 weeks to induce a steatotic and inflammatory milieu (Figure 4A). Mice fed a WD underwent either A/L or an IF (5:2) regimen (Figure 4A). At the end of the 5 weeks, mice fed under A/L conditions were sacrificed in a fed status whilst mice undergoing the IF (5:2) regimen were sacrificed either after a 24-hour fast (fasted state) or following 4 hours of refeeding after a 24-hour fast (refed state) (Figure 4A). By doing so, we aimed to elucidate the key signalling molecules and pathways involved in the fasting response and immediately following refeeding.

**Figure 4.**
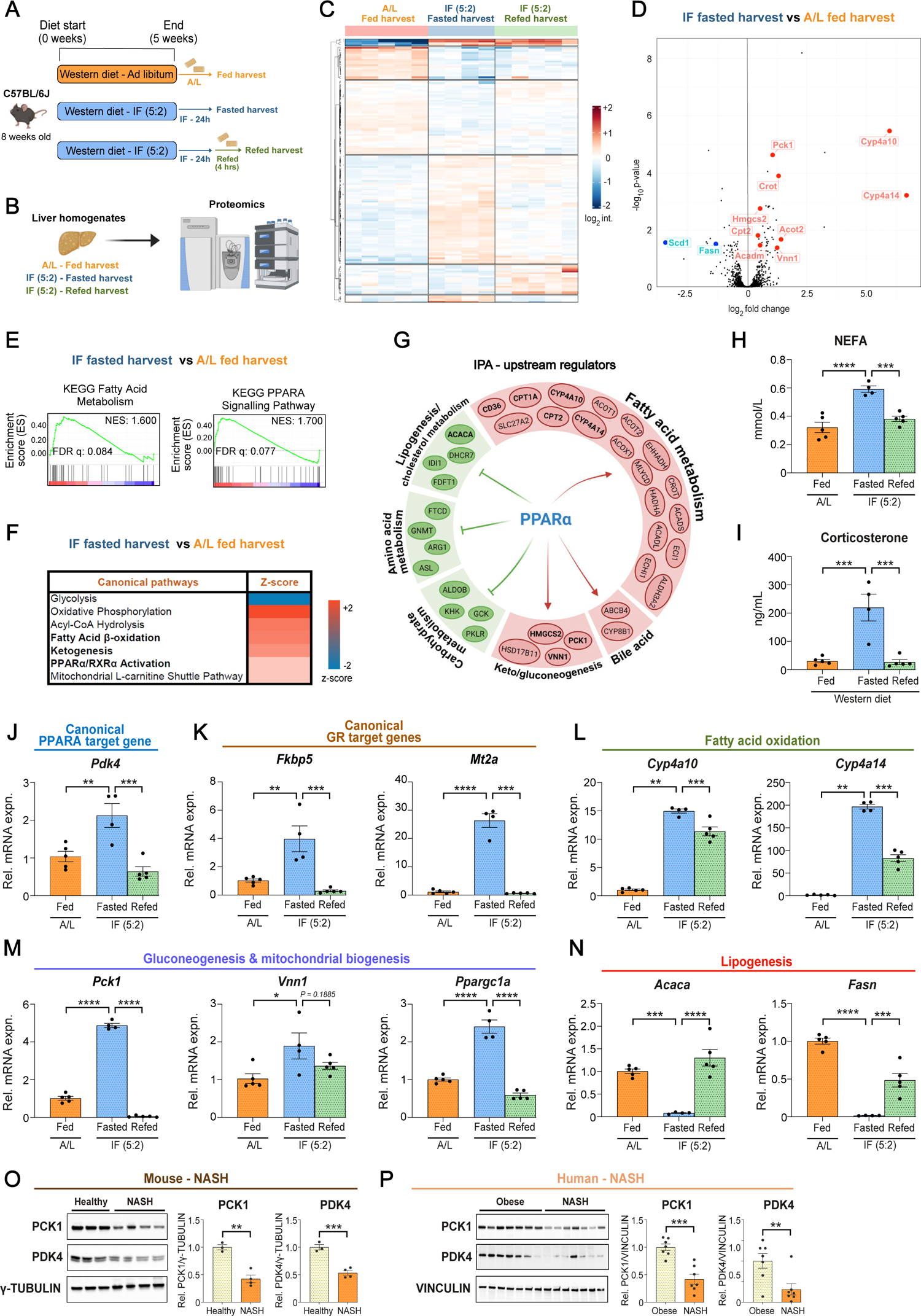
Unbiased proteomics reveal PPARA as a master regulator of the fasting response in NASH. **A.** Scheme of the experiment. 8-week-old C57BL/6J mice were fed a NASH diet for 5 weeks to induce a steatotic and inflammatory milieu. Mice underwent either an *ad libitum* (A/L) or an IF (5:2) regimen. At the end of the 5 weeks, mice fed under A/L conditions were sacrificed in a fed status whilst mice undergoing IF (5:2) regimen were sacrificed either after a 24-hour fast (fasted state) or following 4 hours of refeeding after a 24-hour fast (refed state). **B.** Graphical scheme. Proteomics analysis of the liver homogenates was carried out. **C.** Unsupervised hierarchical clustering revealed distinct groups of proteins regulated under A/L fed (n=5), IF (5:2) fasted (n=4) or IF (5:2) refed (n=5) nutritional states. **D.** Analysis of differentially expressed proteins between IF (5:2) mice (n=4) sacrificed under a fasted state with A/L mice (n=5) sacrificed in a fed state. **E**. Gene Set Enrichment Analysis (GSEA) for ‘KEGG Fatty Acid Metabolism’ and ‘KEGG PPARA Signalling Pathway’ in fasted mice compared to A/L fed controls. **F.** Ingenuity pathway analysis (IPA) revealed ‘Fatty Acid Oxidation’, ‘Ketogenesis’ and ‘PPARα/RXRα Activation’ pathways to be differentially regulated in fasting mice compared to A/L fed controls. **G.** IPA upstream regulator analysis revealed PPARA as a master regulator of the fasting response. **H**. Serum non-esterified fatty acid (NEFA) levels under A/L fed (n=5), IF (5:2) fasted (n=4) or IF (5:2) refed (n=5) nutritional states in mice fed a Western diet (WD) for 5 weeks. **I**. Serum corticosterone levels under A/L fed (n=5), IF (5:2) fasted (n=4) or IF (5:2) refed (n=5) nutritional states in mice fed a WD for 5 weeks. **J-N.** Relative mRNA expression of *Pdk4*, *Fkbp5*, *Mt2a*, *Cyp4a10*, *Cyp4a14*, *Pck1*, *Vnn1, Ppargc1a*, *Acaca* and *Fasn* in liver homogenates of mice fed a WD. mRNA expression was normalized to *Rps14* housekeeping gene. A/L fed (n=5), IF (5:2) fasted (n=4) or IF (5:2) refed (n=5). **O.** Immunoblot images (left) and quantification (right) of PCK1, PDK4 and γ-TUBULIN (loading control) in liver homogenates from healthy (chow-fed; n=3) and NASH (WD-fed; n=4) mice. **P.** Immunoblot images (left) and quantification (right) of PCK1, PDK4 and VINCULIN (loading control) in liver homogenates from obese non-NASH (n=7) and obese NASH (n=7) patients. Data are expressed as mean ± SEM. Statistical significance was calculated using one-way analysis of variance with post-hoc Tukey’s test (H-N) or Student’s t-test (O and P). * P < 0.05, ** P < 0.01, *** P < 0.001, **** P < 0.0001. N.S: non-significant.

We first assessed several key mitogenic and inflammatory signalling pathways involved in NASH pathogenesis and HCC development. Immunoblot analysis revealed that AKT, MTOR-S6K/4EBP1, p38 MAPK and NF-κB pathways were markedly downregulated after a 24-hour fast and increased again following 4 hours of refeeding compared to WD-fed A/L controls (Figure S6A). Given that all of the assessed potentially mitogenic and inflammatory pathways were dampened during the fasting response, suggests that fasting broadly affects multiple pro-growth and inflammatory pathways simultaneously. This could explain, at least in part, some of the benefits of fasting observed in NASH, particularly since each fasting cycle would dampen these pathways in a repetitive manner over the course of a long-term (32 weeks) NASH experiment.

Nevertheless, to better understand the crucial signalling networks orchestrating the fasting response during high-caloric feeding, unbiased proteomics of liver homogenates was performed from A/L fed, IF (5:2) fasted and IF (5:2) refed mice (Figure 4B). Principal component analysis revealed that the IF (5:2) fasted and refed groups clearly cluster separately from A/L fed group (Figure S6B). This was corroborated by unsupervised hierarchical clustering, which revealed distinct groups of proteins regulated under A/L fed, IF (5:2) fasted or IF (5:2) refed nutritional states (Figure 4C). Importantly, analysis of differentially expressed proteins between IF (5:2) mice sacrificed under a fasted state with A/L mice sacrificed in a fed state illustrated that several known fasting response proteins were significantly altered between the two groups (Figure 4D). Specifically, the key gluconeogenic enzymes (*PCK1* and *VNN1),* several enzymes involved in fatty acid oxidation *(CYP4A10, CYP4A14, ACOT2, CROT)* and the rate-limiting enzyme in ketogenesis *(HMGCS2)* were all significantly increased during fasting, whereas key enzymes involved in *de novo* lipogenesis *(FASN* and *SCD1)* were significantly downregulated during fasting (Figure 4D). Moreover, it is also in line with previous observations carried out under normal chow feeding (Hatchwell et al., 2020). This is also in agreement with our whole-body metabolism analyses conducted by indirect calorimetry showing increased fatty acid oxidation and ketogenesis during the fasting response with elevated serum ketone body β-hydroxybutyrate (Figures 2E-2G).

To determine if this fasting phenotype is restricted to a subset of proteins or is more generally applicable to diverse pathways, we performed Gene Set Enrichment Analysis (GSEA) comparing IF (5:2) mice sacrificed under a fasted state with A/L mice sacrificed in a fed state. GSEA revealed a marked enrichment for ‘KEGG Fatty Acid Metabolism’ and ‘KEGG PPARA Signalling Pathway’ and downregulation for ‘HALLMARK MTORC1 Signaling’ and ‘HALLMARK Glycolysis’ signatures in fasted mice compared to A/L fed controls (Figures 4E and S6C). Enrichment for ‘Fatty Acid Oxidation’, ‘Ketogenesis’ and ‘PPARα/RXRα Activation’ was also corroborated by ingenuity pathway analysis (IPA) (Figure 4F). Importantly, search for upstream regulators by IPA revealed peroxisome proliferator-activated receptor alpha (PPARα) to be one of the central mediators of the fasting response during high-caloric feeding (Figure 4G). This is in line with other reports showing that PPARα is a master regulator of the fasting response under chow feeding (Kersten et al., 1999). PPARα resides as a heterodimer with retinoid X receptor alpha (RXRα) and is activated by free fatty acids secreted into the bloodstream via fasting-induced adipose tissue lipolysis (Todisco et al., 2022). Upon binding to free fatty acids, PPARα/RXRα heterodimers recruit co-activators and induce a transcriptional program comprising multiple pathways including fatty acid oxidation (Todisco et al., 2022). Given that PPARα signalling was enriched in the fasted state by proteomics analysis, we analyzed serum non-esterified fatty acid (NEFA) levels under fed, fasted and refed nutritional states in the context of NASH. A significant increase in serum NEFA levels was observed in IF (5:2) mice sacrificed under a fasted state compared to A/L mice sacrificed under a fed state during high-caloric feeding (Figure 4H). Serum NEFA levels rapidly declined following 4 hours of refeeding after a 24-hour fast, illustrating the dynamic nature of the fasting and refeeding process (Figure 4H). Importantly, the fasting response is not solely orchestrated by PPARα but is rather a highly complex process involving dozens of transcription factors at the molecular level but also by multiple hormones at the systemic level (Goldstein et al., 2017; Goldstein and Hager, 2015; Longo et al., 2021). Of these, glucocorticoids (GCs) and the glucocorticoid receptor (GR) signalling have been shown to play a pivotal role in the fasting response as well (Goldstein et al., 2017; Goldstein and Hager, 2015; Præstholm et al., 2020). Recently, it was demonstrated that under normal homeostatic conditions, a concerted effort from both the GR axis and PPARα signalling was required to fully activate the transcriptional program during fasting in the liver (Loft et al., 2022). Therefore, we also assessed whether serum glucocorticoids were elevated during fasting in addition to the increase in serum NEFAs. In line with previous reports (Goldstein et al., 2017), a significant increase in serum corticosterone was found in IF (5:2) mice sacrificed under a fasted state compared to A/L mice sacrificed under a fed state during high-caloric feeding (Figure 4I). Corticosterone is a natural ligand for GRs in hepatocytes (Præstholm et al., 2020). Upon binding to corticosterone, GR translocates to the nucleus and activates a transcriptional program involving fatty acid oxidation, mitochondrial biogenesis and gluconeogenesis (Goldstein et al., 2017; Goldstein and Hager, 2015; Præstholm et al., 2020). Therefore, we next sought to investigate if PPARα and GR target genes are activated during the fasting response. As expected, canonical PPARα and GR target genes including *Pdk4*, *Fkbp5* and *Mt2a* significantly increased during fasting and rapidly declined following refeeding (Figures 4J and 4K). Importantly, we also confirmed our transcriptomics analyses by observing a significant increase in genes involved in fatty acid oxidation, gluconeogenesis and mitochondrial biogenesis in IF (5:2) mice sacrificed under a fasted state compared to A/L mice sacrificed under a fed state (Figures 4L and 4M). Furthermore, we also noted a significant decrease in genes involved in key steps of lipogenesis such as *Acaca* and *Fasn* during fasting, which was reversed upon refeeding (Figure 4N).

Given that the PPARα and GR signalling axes are significantly upregulated during the fasting response, we sought to understand whether these axes, particularly downstream targets of these pathways, are dampened during NASH. Immunoblotting confirmed a significant reduction in the PPARα target gene PDK4 as well as in the GR target gene PCK1 in mouse NASH samples compared to chow controls (Figure 4O). To determine if this applies to humans as well, we analyzed liver samples from obese patients stratified according to their NAS (liver damage severity). By doing so, we could determine if the severity of liver pathology contributes to the regulation of these pathways. Importantly, we confirmed a significant reduction in the expression of PDK4 and PCK1 in human obese patients with NASH (NAS ≥ 5) compared to obese patients without overt liver pathology (NAS = 0) (Figure 4P). Overall, these data clearly demonstrate that the PPARα and GR axes are activated during fasting in response to high-caloric feeding, while these pathways are significantly downregulated in NASH in both mice and men.

### Unbiased metabolomics corroborates fatty acid oxidation and ketogenesis to be highly activated during fasting in NASH

The aberrant metabolic milieu plays an important role in NASH pathogenesis (Ma et al., 2016). Therefore, to determine whether IF also influences the metabolome in the context of chronic high-caloric feeding, we performed unbiased metabolomics (by quantitative NMR spectroscopy) of liver homogenates from A/L fed, IF (5:2) fasted as well as IF (5:2) refed mice as before (Figure 5A). Similar to proteomics, principal component analysis revealed that all three groups cluster separately, with the largest separation caused by the IF (5:2) fasted group (Figure 5B). This was corroborated by unsupervised hierarchical clustering, which revealed a total of 20 polar metabolites (out of 57 compounds) to be significantly differentially regulated under the different nutritional states (Figure 5C). Further corroborating the proteomics analyses and the indirect calorimetry, a significant increase in the liver ketone body 3-hydroxybutyrate and a decrease in glucose and lactate were found during fasting, which was mostly reversed during refeeding, indicating elevated ketogenesis and a dampening of glycolysis in the fasted state (Figures 5D and 5E). Elevated levels of AMP and GMP were also found during fasting, indicating a state of low energy (Figure 5E). Interestingly, increased levels of antioxidant-related compounds such as taurine, glutathione, NADP+ and ß-alanine were found in the IF (5:2) mice during fasting and/or refeeding, which suggests that antioxidant mechanisms are initiated during the fasting and refeeding cycles (Figure 5F). Of note, there was a remarkable increase in glutathione only in the refed group. Glucose levels in the refed animals did not reach the same levels as in the fed regimen, which could be a result of increased glucose uptake in cells induced by ß-alanine (Billacura et al., 2022). Pathway analysis to assess this process in more depth confirmed that pathways for ‘glycolysis’, ‘Warburg effect’ and ‘pyruvate metabolism’ were all significantly dampened during fasting, whilst there was a significant enrichment for ‘fatty acid metabolism’, ‘bile acid homeostasis’ and ‘taurine/hypotaurine metabolism’ (Figure 5G). Conversely, in the refed state, many of these processes were reversed such as the ‘Warburg effect’, ‘pyruvate metabolism’ and ‘fatty acid metabolism’ (Figure 5H). Overall, these data illustrate that fasting and refeeding cycles induce a profound metabolic shift promoting fatty acid oxidation, ketogenesis, increased cellular glucose uptake and antioxidant processes, which together act as beneficial detoxification mechanisms that may help to combat NASH.

**Figure 5.**
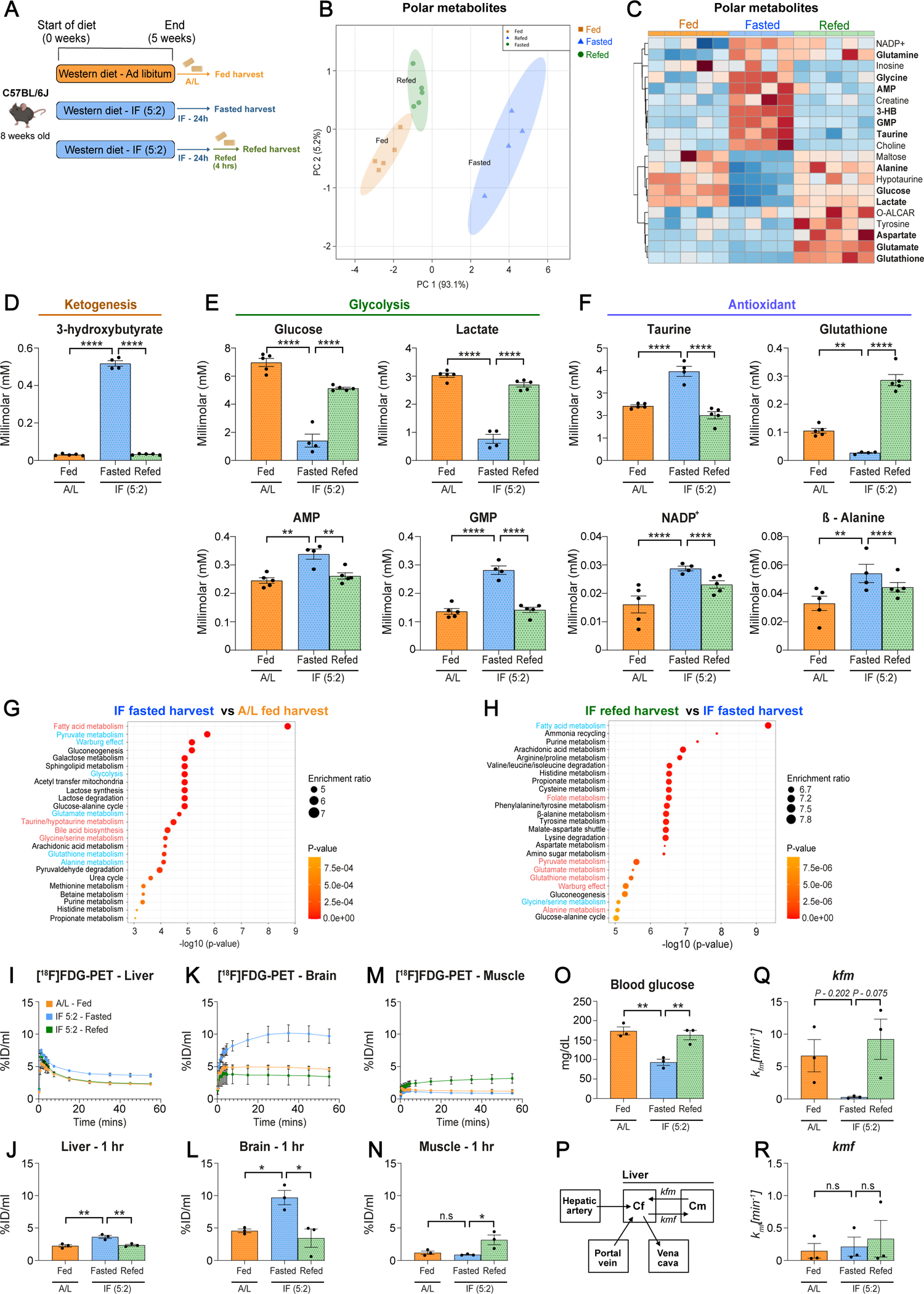
Unbiased metabolomics reveal fatty acid oxidation and ketogenesis are activated during fasting in NASH. **A.** Experimental scheme. 8-week-old C57BL/6J mice were fed either a chow diet or a NASH-inducing diet for 5 weeks in the presence or absence of IF 5:2 fasting regimen where mice underwent two fasting cycles per week on two non-consecutive days with each fasting cycle lasting for a duration of 24 hours. At the end of the 5 weeks, mice fed under A/L conditions were sacrificed in a fed status whilst mice undergoing IF (5:2) regimen were sacrificed either after a 24-hour fast (fasted state) or following 4 hours of refeeding after a 24-hour fast (refed state). **B.** PCA of the three-group comparison illustrating Western Diet control (orange), fasted (blue) and refed (green) group 95% confidence intervals and group separation based on different polar metabolite concentrations. **C.** Heatmap generated by unsupervised clustering of NMR spectroscopy-based metabolomics analysis of livers illustrating the 20 ANOVA statistically significant metabolites (red – increased concentration, blue – reduced concentration). **D-F.** Key metabolic pathways, such as ketogenesis, glycolysis and antioxidant balance, upregulated or downregulated in the context of fasting and refeeding stimuli, illustrated as individual metabolite concentration bar plots. **G and H.** SMPDB library-based pathway enrichment analysis for significantly regulated pathways in different nutritional state such as (g) fasted vs. fed and (h) refed vs. fasted. **I-R.** Dynamic [^18^F]FDG-PET scans were performed from A/L fed, IF (5:2) fasted as well as IF (5:2) refed mice on WD for 32 weeks. The IF (5:2) regimen was carried out for the final 12 weeks. At this point, [^18^F]FDG was injected into A/L fed mice or in mice undergoing the IF (5:2) regimen after a 24-hour fast (fasted state) or following 4 hours of refeeding after a 24-hour fast (refed state). **I-N.** Dynamic [^18^F]FDG-PET scans (top) and the corresponding quantifications at the 1 hour time point (bottom) of the indicated groups in the liver, brain and skeletal muscle. **O.** Blood glucose levels between WD A/L-fed and IF (5:2) fasted/refed mice prior to injection of [^18^F]FDG. **P.** Schematic illustrating the compartmental modelling approach using the gut tracer concentration to calculate the input function of the portal vein. **Q.** The rate constant *k_fm_* representing the transport of [^18^F]FDG from the metabolized/phosphorylated compartment (Cm) to the free compartment (Cf) in the liver between the indicated groups. **R.** The reverse rate constant *k_mf_* (Cf to Cm) of the indicated groups. All data are shown as mean ± SEM. Statistical significance was calculated using parametric ANOVA and Fischer’s least significant difference (LSD) test. ** P < 0.01, *** P < 0.001, **** P < 0.0001. N.S: non-significant.

### Dynamic [^18^F]FDG-PET *in vivo* revealed the adaptions of glucose metabolism induced by fasting

To assess the impact of fasting on the *in vivo* liver glucose metabolism, dynamic [^18^F]FDG-PET scans were performed from A/L fed, IF (5:2) fasted as well as IF (5:2) refed mice on WD for 32 weeks. The IF (5:2) regimen was carried out for the final 12 weeks. At this point, [^18^F]FDG was injected into A/L fed mice or in mice undergoing the IF (5:2) regimen after a 24-hour fast (fasted state) or following 4 hours of refeeding after a 24-hour fast (refed state).

Time activity curves (TAC) revealed a significantly enhanced [^18^F]FDG uptake in the liver and to a greater extent in the brain after 24 hours of fasting in mice compared to A/L fed controls (Figures 5I-5L). This is in line with the requirement of glucose as a fuel source for the brain following the use of ketone bodies during a fasting state. The increase in glucose uptake was already compensated after 4 hours of refeeding, corroborated by proteomics and metabolomics data showing the dynamic nature of the fasting and refeeding cycles. In contrast, [^18^F]FDG uptake in muscle was significantly higher following 4 hours of refeeding after a 24-hour fast (Figures 5M and N). Importantly, the enhanced uptake of [^18^F]FDG in the refed state was not simply due to differences in blood glucose levels, as the blood glucose levels were comparable between WD A/L-fed and IF (5:2) refed mice prior to injection of [^18^F]FDG (Figure 5O). This implies enhanced insulin-induced glucose uptake in skeletal muscle in mice undergoing the IF (5:2) regimen, which is in accordance with previous reports showing that IF can improve glucose homeostasis and insulin sensitivity even in humans (Sutton et al., 2018).

The quantification of hepatic glucose metabolism by [^18^F]FDG-PET is more complex than in other tissues, as hepatocytes are able to phosphorylate and dephosphorylate [^18^F]FDG/glucose by glucose-6 phosphatase activity. Furthermore, the tracer is distributed to the liver by the dual blood supply of the hepatic artery and the portal vein (Figure 5P). Therefore, we facilitated a compartmental modelling approach developed by (Garbarino et al., 2015), which is considering this special situation by using the gut tracer concentration to calculate the input function of the portal vein (Figure 5P). The rate constant *k_fm_* representing the exchange rate of [^18^F]FDG from the metabolized/phosphorylated compartment (Cm) to the free compartment (Cf) in the liver was lower in IF (5:2) mice after a 24 hour fast compared to A/L fed controls (Figure 5Q). The reverse rate constant *k_mf_* (Cf to Cm) was not significantly altered between groups (Figure 5R).

These data suggest that the 24h-fasting periods did influence the hepatic glucose metabolism, particularly by inhibiting the efflux of phosphorylated [^18^F]FDG out of the hepatocytes. We did not observe a downregulation of glucose 6-phosphatase in the proteomics analyses, and this is in line with activation of gluconeogenesis via PPARα and/or PCK-1 by the fasting regimen (Figure 4). Furthermore, the blood glucose levels were significantly lower in the IF groups, therefore higher insulin levels can also be excluded to interfere with glucose 6-phosphatase expression and the decrease in *k_fm_* (Figure 5O). Most likely, [^18^F]FDG needs to compete with the amount of glucose generated by gluconeogenesis for the phosphatase activity of the enzyme glucose 6-phosphatase. This may lead to lower efflux rates of [^18^F]FDG (*k_fm_*) from the hepatocytes to the free compartment compared to the unfasted animals. Therefore, compartmental modelling of [^18^F]FDG-PET data might be a non-invasive method to asses the *in vivo* gluconeogenesis induction in the liver by a fasting regimen.

### Hepatocyte-specific deletion of *GR* does not fully abrogate the hepatic fasting response

As we observed a significant increase in serum GCs and GR signalling targets in the liver during fasting, we sought to understand if GC-GR signalling plays a critical role in the hepatic fasting response (Figure S7A). Therefore, we fed hepatocyte-specific GR knockout mice (*Alb-cre x GR^fl/fl^ – GR^HEP^*) and control littermates (*GR^fl/fl^*) a WD for 5 weeks under both A/L or IF (5:2) conditions and sacrificed the mice in either a fed or fasted state, respectively (Figure S7B). Fasted mice showed lower blood glucose levels and increased serum ketone body β-hydroxybutyrate irrespective of the genotype (Figure S7C), indicating that hepatocyte-glucocorticoid signalling does not play a major role in fasting-induced ketone body production in the context of WD feeding. Nevertheless, fasting-induced hepatic expression of canonical GR target genes *Fkbp5* and *Mt2a* was abrogated in *GR^HEP^* mice (Figure S7D). Additionally, fasting-induced expression of fatty acid oxidation genes (e.g., *Cyp4a10*) in the liver was significantly impaired, but not completely abrogated in *GR^HEP^* mice (Figure S7E). In contrast, fasting-induced hepatic expression of the key mitochondrial biogenesis gene *Ppargc1a* and the rate-limiting enzyme in gluconeogenesis *Pck1* was entirely prevented in *GR^HEP^* mice (Figures S7F and S7G). *GR^HEP^*mice did not show any alterations in fasting-induced suppression of anabolic lipogenesis (Figure S7H). Overall, these data demonstrate that hepatocyte GR-GC signalling regulates several aspects of the hepatic fasting response but is not solely responsible for orchestrating this response, particularly in the regulation of fasting-induced fatty acid oxidation. This would be in line with the proteomics analyses alluding to the role of PPARα to be a central player in the hepatic fasting response.

### Combined depletion of *Ppara* and *Pck1* markedly dampens the fasting response

As aforementioned, the hepatic fasting response under a basal (chow feeding) state is mediated by a concerted effort of many hormonal and transcriptional networks (Goldstein et al., 2017; Goldstein and Hager, 2015; Longo et al., 2021). In agreement, we have observed an elevation of NEFAs and GCs in the serum of fasted mice in NASH with a subsequent activation of hepatic PPARα and GR signalling, confirmed by induction in the expression of multiple genes involved in the fasting response including *Cyp4a10* and *Pck1* (Figures 4G, 4H, and 6A). We then aimed to understand if these axes truly play a causal role in mediating the hepatic fasting response in NASH or are merely correlative. Therefore, we depleted *Ppara* or *Pck1* in isolation or both together using adeno-associated virus vectors (AAV8) in C57BL/6J mice (Figure 6B). We then fed these mice a WD under A/L conditions or with an IF (5:2) regimen for 5 weeks, and as above, sacrificed the mice in either a fed or fasted state (Figures 6B and 6C).

**Figure 6.**
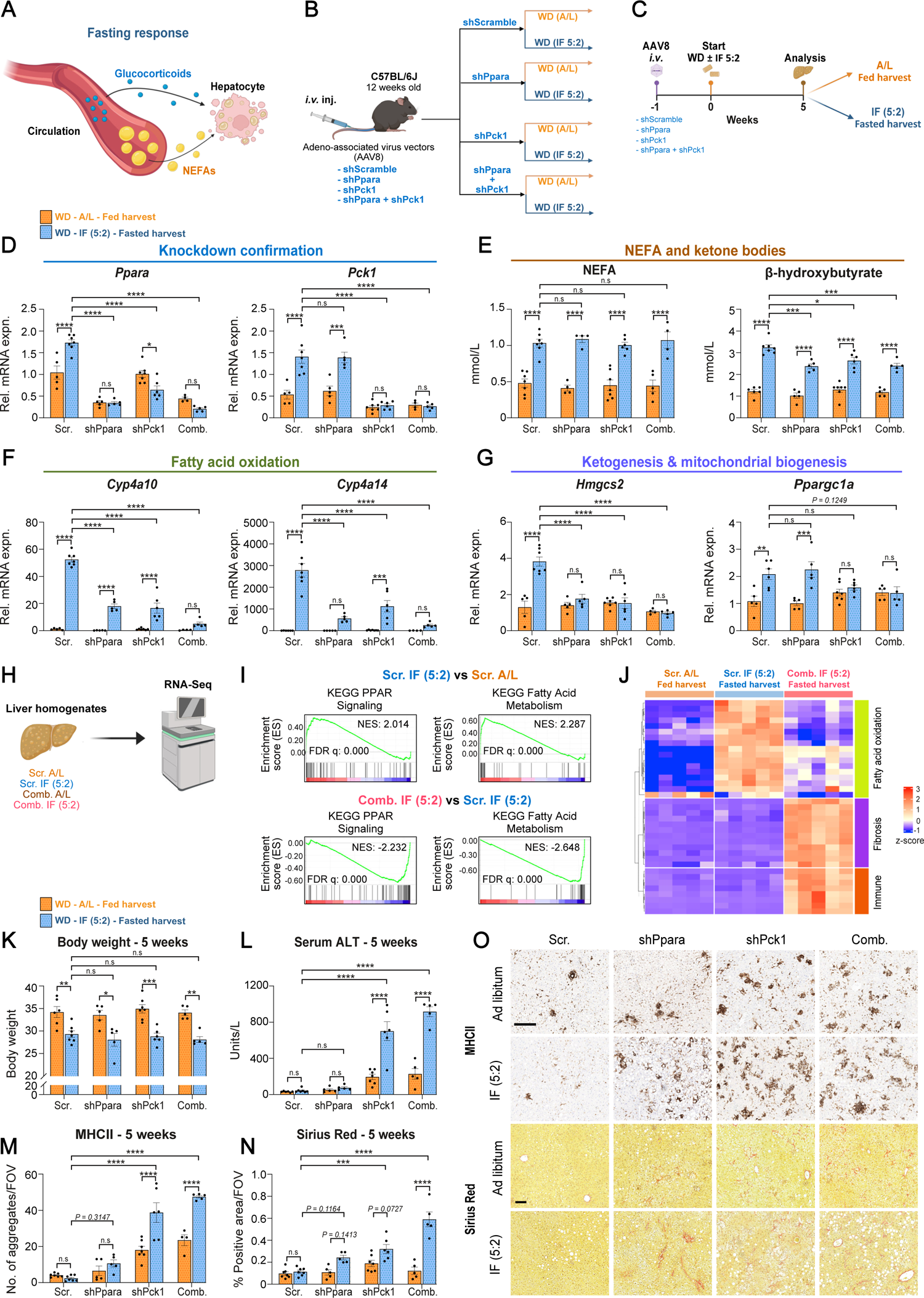
PPARA and glucocorticoid-PCK1 axis are critical for mediating the benefits of intermittent fasting in NASH. **A.** Graphical scheme. Fasting induces an increase in serum non-esterified fatty acids (NEFAs) and glucocorticoids (GCs), which in turn activate PPARA and glucocorticoid receptor (GR)-PCK1 signalling in hepatocytes, respectively. **B and C.** Scheme of AAV8-mediated knockdown of *Ppara* and *Pck1* alone or both together. 12-week-old C57BL/6J mice were injected intravenously (*i.v.)* with AAV8 vectors carrying either a scrambled shRNA or shRNAs against *Ppara* or *Pck1*. Mice were then divided into two groups with one group being fed a Western diet (WD) for 5 weeks under *ad libitum* (A/L) conditions whilst the other group underwent an IF (5:2) regimen. At the end of the 5 weeks, mice fed under A/L conditions were sacrificed in a fed status whilst mice undergoing IF (5:2) regimen were sacrificed after a 24-hour fast (fasted state). **D.** Relative mRNA expression of *Ppara* and *Pck1* in liver homogenates of the indicated genotypes in mice fed a WD. mRNA expression was normalized to *Rps14* housekeeping gene. Scr. A/L (n=5), Scr. IF (5:2) (n=7), shPpara A/L (n=5), shPpara IF (5:2) (n=5), shPck1 A/L (n=7), shPck1 IF (5:2) (n=6), Comb. A/L (n=4) and Comb. IF (5:2) (n=5). **E.** Serum non-esterified fatty acid (NEFA) or β-hydroxybutyrate levels of the indicated genotypes in mice fed a WD. **F and G.** Relative mRNA expression of *Cyp4a10, Cyp4a14, Hmgcs2* and *Ppargc1a* of the indicated genotypes in mice fed a WD. mRNA expression was normalized to *Rps14* housekeeping gene. Scr. A/L (n=5-6), Scr. IF (5:2) (n=6-7), shPpara A/L (n=5), shPpara IF (5:2) (n=5), shPck1 A/L (n=7), shPck1 IF (5:2) (n=6), Comb. A/L (n=4-5) and Comb. IF (5:2) (n=5). **H.** Graphical scheme. Bulk RNA-sequencing of whole liver lysates from the indicated groups were carried out. **I**. Gene Set Enrichment Analysis (GSEA) for ‘KEGG PPARA Signalling Pathway’, ‘KEGG Fatty Acid Metabolism’, ‘REACTOME Glycolysis’, REACTOME Cholesterol Biosynthesis’. ‘HALLMARK Inflammatory Response’ and ‘REACTOME Collagen Formation’ of the indicated groups. **J.** Heatmap of differentially expressed genes involved in fatty acid oxidation, fibrosis and inflammation of the indicated groups. **K and L.** Body weight and serum ALT at the end-point (5 weeks of WD) from the indicated. Scr. A/L (n=6), Scr. IF (5:2) (n=7), shPpara A/L (n=5), shPpara IF (5:2) (n=5), shPck1 A/L (n=7), shPck1 IF (5:2) (n=6), Comb. A/L (n=5) and Comb. IF (5:2) (n=5). **M and N.** Quantification of MHCII aggregates and Sirius Red staining of the indicated genotypes in mice fed a WD. Scr. A/L (n=7), Scr. IF (5:2) (n=7), shPpara A/L (n=5), shPpara IF (5:2) (n=5), shPck1 A/L (n=7), shPck1 IF (5:2) (n=6), Comb. A/L (n=4-5) and Comb. IF (5:2) (n=5). **O.** Representative MHCII (top) and Sirius Red (bottom) immunohistochemistry images of the indicated groups at the end-point (5 weeks of WD). Data are expressed as mean ± SEM. Statistical significance was calculated using two-way analysis of variance with post-hoc Tukey’s test. * P < 0.05, ** P < 0.01, *** P < 0.001, **** P < 0.0001. N.S: non-significant. Scale bar, 100µm.

We first confirmed knockdown of *Ppara* and/or *Pck1* under both A/L and fasted conditions (Figure 6D). Surprisingly, we observed that *Ppara* expression is downregulated upon depletion of *Pck1* under a state of fasting but not during a fed harvest (Figure 6D). In contrast, depletion of *Ppara* did not affect the expression of *Pck1* in either fed or fasted states (Figure 6D). As before, fasted mice showed a significant reduction in blood glucose levels and a concomitant increase in serum NEFAs compared to A/L fed controls irrespective of the genotype, which clearly indicates that the systemic fasting response is unaltered upon knockdown of *Ppara* and/or *Pck1* (Figures 6E and S8A). Nevertheless, the fasting-induced elevation of serum ketone body β-hydroxybutyrate was significantly reduced upon depletion of *Ppara* and/or *Pck1* (Figure 6E). Additionally, expression of genes involved in fasting-induced fatty acid oxidation and ketogenesis was significantly impaired upon depletion of *Ppara* or *Pck1* with the greatest effect with combined depletion (Figures 5F, 5G, and S8B). This corroborates previous findings showing that hepatocyte-specific deletion of *Ppara* abrogates fasting-induced upregulation of key fatty acid (*e.g., Cyp4a14*) and ketogenic genes (*e.g., Hmgcs2*) under chow-feeding (Montagner et al., 2016). Fasting-induced expression of the key mitochondrial biogenesis enzyme *Ppargc1a* was prevented only upon depletion of *Pck1,* indicating that there are subsets of genes differentially regulated by PPARα and PCK1 (Figure 6G). Conversely, fasting-induced upregulation of the key hepatic gluconeogenic enzyme *Vnn1* (the other being *Pck1*) was only prevented upon depletion of *Ppara* but not *Pck1* (Figure S8C). Interestingly, fasting-induced suppression of *de novo* lipogenesis (*Fasn*) was not affected by depletion of *Ppara* and/or *Pck1* (Figure S8D).

We then investigated whether depletion of *Ppara* and/or *Pck1* affected the fasting response in the liver more broadly. To tackle this, we carried out RNA-sequencing of livers from all of the distinct groups. We initially focused on the effect of combined depletion of *Ppara* and *Pck1* on the fasting response compared to scramble A/L controls (Figure 6H). PCA revealed a clear separation between the groups (Scr. A/L, Scr. IF (5:2), Comb. A/L and Comb. IF (5:2)) (Fig. S8E). In accordance with the proteomics analysis, GSEA of transcriptomics data revealed a significant enrichment for ‘KEGG PPAR Signaling’ and ‘KEGG Fatty Acid Metabolism’ in Scr. IF (5:2) mice compared to Scr. A/L controls (Figures 6I and 6J). Importantly, the combined knockdown of *Ppara* and *Pck1* abrogated the fasting-induced enrichment of these selected pathways (Figures 6I and 6J). Surprisingly, the combined knockdown of *Ppara* and *Pck1* in the context of fasting showed a significant enrichment for pathways related to inflammation and fibrosis when compared to Scr. IF (5:2) mice (Figures 6J and S8F). Importantly, depletion of *Ppara* or *Pck1* alone in the context of fasting also showed a transcriptome profile very similar to combined depletion with downregulation of pathways related to fatty acid metabolism and a concomitant enrichment for inflammatory and fibrosis signatures (Figures S8G-S8I). Altogether, these data suggest that depletion of *Ppara* and/or *Pck1* not only abrogates the transcriptional fasting response but rather also exacerbates liver pathology. This highlights the crucial role PPARα and/or PCK1 as well as their downstream targets play in mediating the hepatic fasting response and enabling liver homeostasis. Therefore, the lack of these proteins upon fasting may be detrimental, particularly in the context of NASH.

To determine if this is truly the case, we then assessed the effects of knockdown of *Ppara* and/or *Pck1* on systemic obesity and liver pathology. As before, fasted mice displayed a reduction in body weight compared to A/L fed controls irrespective of the genotype (Figures 6K, S8J, and S8K). This further highlights that depletion of *Ppara* and/or *Pck1* does not affect systemic obesity or metabolism (Figure 6K). Conversely, serum ALT, a marker of liver damage, was drastically increased in mice with knockdown of *Pck1* or combined depletion of *Ppara* and *Pck1* under A/L conditions but this was exacerbated in mice undergoing IF (5:2) regimen (Figure 6L). In support, depletion of *Ppara* and/or *Pck1* in the context of fasting exacerbated WD-induced hepatomegaly (Figures S8L and S8M). This suggests that knockdown of key executors of the hepatic fasting response not only abrogates the benefits of fasting in NASH but rather is actively detrimental. We therefore interrogated this in more detail at histopathological level. In support, depletion of *Pck1* and to a lesser extent *Ppara* in the context of fasting significantly aggravated WD-induced immune infiltration and fibrosis, which corroborated transcriptomic analyses (Figures 6M-6O, S8N, and S8O). Overall, *Ppara* and *Pck1* act as hepatic executors of the fasting response and knockdown of these targets not only abrogates the hepatic transcriptional program induced by fasting but also significantly worsens liver pathology without affecting systemic obesity or metabolic parameters.

### PPARα agonist pemafibrate recapitulates aspects of the hepatic fasting response

Given that PPARα was at the center of the hepatic fasting response in our proteomic and transcriptomic analyses and that pharmacological agents are available to target this axis, we aimed to understand if the PPARα agonist pemafibrate could mimic fasting-induced molecular pathways in the liver (Figure S9A). Of note, there are currently no available pharmacological agents to agonize PCK1. We therefore, treated 8-week-old C57BL/6J mice with either pemafibrate or vehicle for 24 hours and compared this to mice that were fasted for 24 hours (Figure S9B). Whilst fasting lowered body weight and blood glucose levels, pemafibrate had no effect on these systemic parameters (Figure S9C). In contrast, pemafibrate induced expression of the PPARα target gene *Pdk4* to a much greater extent than fasting (Figure S9D). Pemafibrate also induced genes involved in fatty acid oxidation (e.g., *Cyp4a10*) but not to the same level as fasting (Figure S9E). Pemafibrate, however, induced the key ketogenic enzyme *Hmgcs2* and the gluconeogenic enzyme *Vnn1* to the same degree as fasting (Figure S9F). Although fasting significantly suppressed anabolic lipogenesis, pemafibrate had no effect on the expression of these enzymes (Figure S9G). Overall, these data indicate that whilst PPARα agonism alone mimics some aspects of the hepatic fasting response, it is not sufficient to fully recapitulate this process, in particular the induction of fatty acid catabolism whilst suppressing anabolic lipogenesis. This is in line with our data that shows it is a concerted effort of both PPARα and GR-PCK1 action that mediates the hepatic fasting response.

### Therapeutic fasting ameliorates established NASH as well as blunts subsequent liver cancer transition in two distinct obesogenic models of NASH-HCC

Finally, we investigated the therapeutic potential of the IF (5:2) regimen in established NASH. Therefore, we first fed 8-week-old male C57BL/6J mice with WD for 5 months to induce NASH (Figure 7A). At this point, mice were segregated into two groups matched for their body weight (Figure S10A). One group was given A/L access to food and water whilst the other group underwent the IF (5:2) regimen carried out in the active phase (7pm to 7pm) (Figure 7A). Mice were continued on the WD for 4 further months (total of 9 months). Chow-fed mice (ND) served as controls (Figure 7A).

**Figure 7.**
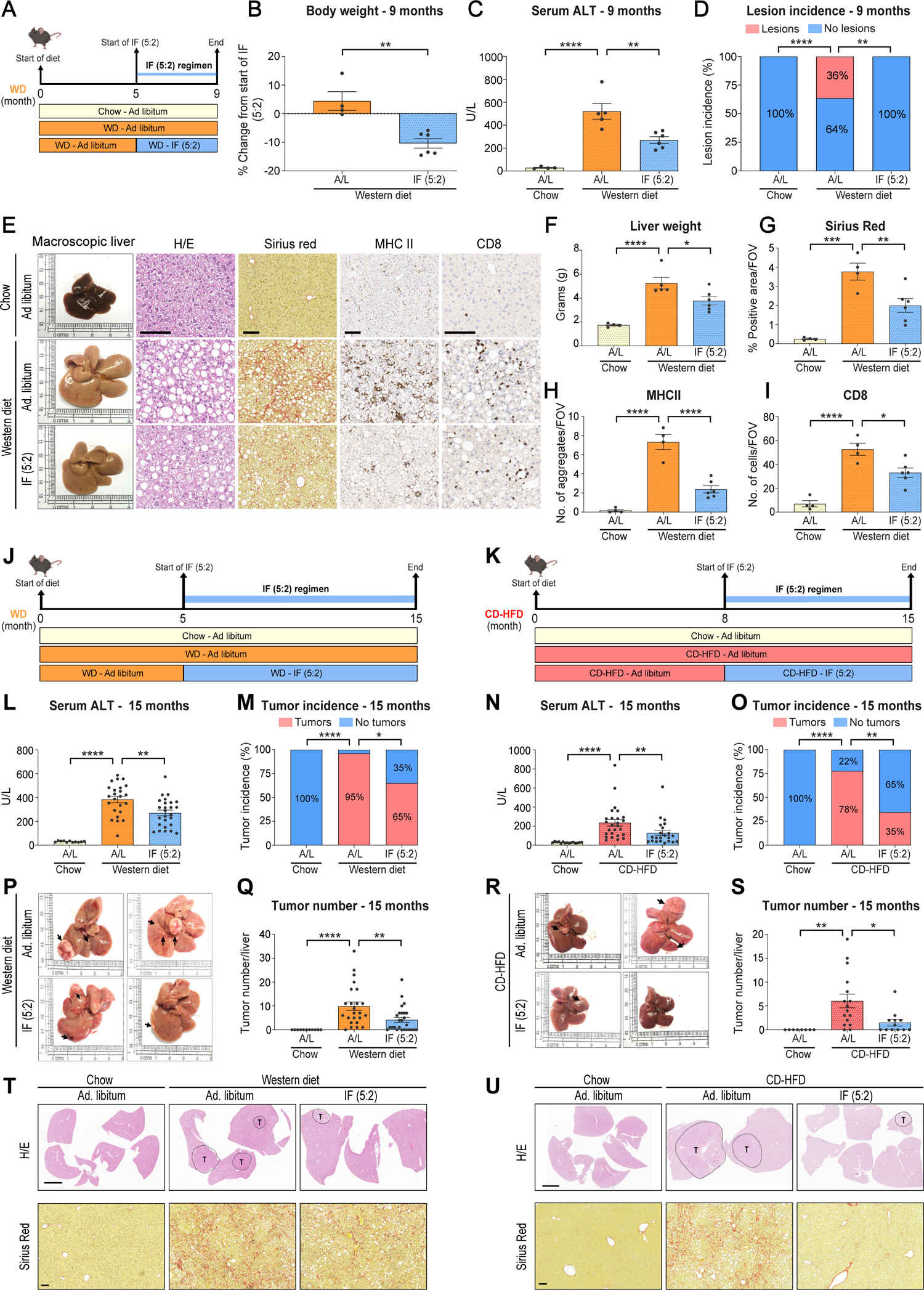
Therapeutic intermittent fasting reverts established NASH and diminishes subsequent transition to hepatocellular carcinoma. **A.** Experimental scheme. 8-week-old male C57BL/6J mice were fed either normal chow diet or a Western diet (WD) for 5 months to induce NASH. At this point, mice were evenly distributed to the different treatment arms. Mice were then continued on the diets under *ad libitum* (A/L) conditions or underwent the IF (5:2) regimen for 4 additional months to assess NASH amelioration. **B.** Body weight change following 4 months of IF 5:2 regimen. WD A/L (n=4) and WD IF (5:2) (n=6). **C.** Serum alanine aminotransferase (ALT) at the end-point (9 months of WD). Chow A/L (n=4) WD A/L (n=5) and WD IF (5:2) (n=6). **D.** Lesion incidence (%) at the end-point (9 months of WD). Chow A/L (n=8) WD A/L (n=11) and WD IF (5:2) (n=11). **E.** Representative macroscopic liver photos and immunohistochemistry images for H/E, Sirius Red, MHCII and CD8 staining at the end-point (9 months of WD). **F.** Absolute liver weight. Chow A/L (n=4) WD A/L (n=5) and WD IF (5:2) (n=6). **G-I.** Quantification of Sirius Red, MHCII and CD8 stainings. Chow A/L (n=4) WD A/L (n=4) and WD IF (5:2) (n=6). **J and K.** Scheme of the experiment. 8-week-old male C57BL/6J mice were fed with either a WD for 5 months or choline-deficient high-fat diet (CD-HFD) for 8 months to induce NASH. At this point, mice were evenly distributed to the different treatment arms. Mice were then continued on the diets under A/L conditions or underwent the IF (5:2) regimen to assess NASH-HCC transition 15 months post-diet start. **L and N.** Serum ALT at the end-point (15 months of WD or CD-HFD) under A/L or IF (5:2) regimen. Chow A/L (11), WD A/L (n=24) and WD IF (5:2) (n=26). Chow A/L (n=15), CD-HFD A/L (n=26) and CD-HFD IF (5:2) (n=22). **M and O.** Liver cancer incidence (%) at the end-point (15 months of WD or CD-HFD) under A/L or IF (5:2) regimen. Chow A/L (n=11), WD A/L (n=24) and WD IF (5:2) (n=26). Chow A/L (12), CD-HFD A/L (n=27) and CD-HFD IF (5:2) (n=23). **P and R.** Representative macroscopic liver photos at the end-point (15 months of WD or CD-HFD) IF (5:2) regimen. Arrows indicate tumor nodules. **Q and S.** Liver tumor numbers at the end-point (15 months of WD or CD-HFD) under A/L or IF (5:2) regimen. Chow A/L (11), WD A/L (n=24) and WD IF (5:2) (n=26). Chow A/L (n=8), CD-HFD A/L (n=16) and CD-HFD IF (5:2) (n=26). **T and U.** Representative immunohistochemistry images for H/E and Sirius Red at the end-point (15 months of WD or CD-HFD). Data are expressed as mean ± SEM. Statistical significance was calculated using Student’s t-test (B), one-way analysis of variance with post-hoc Tukey’s test (C,F-I,L,N,Q,S) or Fisher’s exact test (d.m and o). (* P < 0.05, ** P < 0.01, *** P < 0.001, **** P < 0.0001). N.S: non-significant. Scale bar, 100µm for E. Scale bar, 5mm for T (H/E) and 100µm T (Sirius Red).

Importantly, the IF (5:2) regimen carried out in a therapeutic manner not only induced a significant reduction in body weight and fat mass but also in serum ALT, ALP and TCHOL levels compared to WD-fed A/L controls (Figures 7 B, 7C, and S10B-S10E). Additionally, a subset (36%) of WD-fed A/L mice displayed early neoplastic lesions whilst none of the fasted mice showed lesions (Figure 7D). As before, WD-fed A/L controls presented with overt hepatomegaly, hepatic steatosis, fibrosis and immune infiltration compared to ND-fed controls (Figures 7E-7I and S10F). Strikingly, the IF (5:2) regimen even when carried out in a therapeutic manner displayed significantly lower liver weight (g and %), fibrosis as well as myeloid (MHCII) and lymphoid (CD8) cell infiltration (Figures 7E-7I and S10F).

Furthermore, we tested whether the IF (5:2) regimen can ameliorate NASH in the CD-HFD model with the same experimental setting as above (Figure S10G). As before, following 5 months of CD-HFD feeding to induce NASH, mice were segregated into two groups matched for their body weight (Figure S10H). One group of mice was then subjected to the IF (5:2) regimen for 4 further months (total of 9 months) (Figure S10G). We observed a significant reduction in body weight, serum liver damage markers (ALT and ALP) as well serum cholesterol in mice subjected to IF (5:2) compared to A/L CD-HFD controls (Figures S10I-10N). Furthermore, mice undergoing IF (5:2) also showed a significantly lower liver weight, hepatic steatosis, fibrosis as well as myeloid (MHCII) and lymphoid (CD8) cell infiltration (Figures S10O-S10S). Overall, this clearly demonstrates that the IF (5:2) regimen ameliorates established NASH in multiple obesogenic models of NASH and highlights that this could be a viable intervention against NASH in patients.

Upon observing that the IF (5:2) regimen can ameliorate established NASH, we then investigated if this regimen can also blunt NASH to HCC transition when carried out in a therapeutic manner. To tackle this comprehensively, we once again utilized both the WD and CD-HFD models of NASH and NASH-HCC (Figures 7J and 7K). Therefore, C57BL/6J mice were first fed either WD (5 months) or CD-HFD (8 months) to induce NASH (Figures 7J and 7K). At this point, mice were segregated into two groups matched for their body weight and serum ALT values (Figures S11A-11D). Mice were then continued under A/L access to food and water or underwent the IF (5:2) regimen carried out in the active phase (7pm to 7pm) (Figures 7J and 7K). Mice were fed a WD or CD-HFD for a total of 15 months. Chow-fed mice (ND) served as controls (Figures 7J and 7K). Notably, given the length of these experiments (15 months), it is imperative to highlight that our aim was to study true spontaneous NASH-HCC from feeding alone to closely mimic the human pathophysiology as opposed to the use of carcinogens (e.g., DEN) or hepatotoxins (CCl_4_) to accelerate the process.

Mice undergoing therapeutic IF (5:2) displayed significantly lower body weight, fat mass, and serum ALT and TCHOL values compared to A/L controls in both the WD and CD-HFD models of NASH-HCC (Figures 7L, 7N, and S11E-S11L). Importantly, fasted mice not only showed significantly lower liver tumor incidence but strikingly lower tumor numbers as well compared to A/L controls in both the WD and CD-HFD models of NASH-HCC (Figures 7M, 7O, and 7P-7U). Nevertheless, no significant difference in tumor quality was detected between A/L controls and fasted mice (Figure S12). Fasted mice, however, displayed significantly lower fibrosis and immune cell infiltration (Figures 7T and U, and fig. S13, M to V). Overall, the IF (5:2) regimen not only ameliorated NASH when applied in a therapeutic manner but also significantly blunted the subsequent development of HCC in two distinct diet-induced models of NASH-HCC. Therefore, the IF (5:2) regimen could hold great therapeutic value as a potent and effective intervention for NASH.

## Discussion

Dietary and fasting regimens have been religiously followed for millennia and the scientific community has increasingly begun to recognize their benefits in obesity, metabolic diseases and even cancer (Taylor et al., 2022). Our findings clearly demonstrate that the IF 5:2 regimen is effective, when applied both as a preventive or therapeutic intervention, against the development and treatment of NASH and fibrosis. Similarly, the IF 5:2 regimen also significantly blunted the development of NASH-HCC. Importantly, not all fasting regimens evoked the same beneficial response against NASH since the timing, length and number of fasting cycles as well as the type of NASH diet all contributed to the effectiveness of the fasting regimen; thus highlighting that there are different ‘flavors’ to these regimens. For example, fasting in the active phase was more effective than in the inactive phase. Importantly, serum corticosterone in mice is at its highest around 6:30 pm prior to the onset of their active phase (Kakihana and Moore, 1976). Given the importance of glucocorticoid signalling in mediating the hepatic fasting response, this may explain why initiating the fasting response in the active phase is more beneficial than when initiated in the inactive phase. Others have also shown that dietary regimens such as TRF can reset the circadian rhythm that is disrupted in obesity and metabolic diseases (Chaix et al., 2019). Additionally, the number of fasting cycles needed per week to obtain the benefits was dependent on the aggressiveness of the NASH diet. This would be in line with a more rigorous fasting regimen (IF 5:2 > IF 6:1) for patients with more severe NASH. The IF 5:2 regimen should be more compliance-friendly than undergoing long-term calorie deficits since patients can still consume not only the same quantity of food but the type of foods that they want as well in the feeding days, as it is notoriously difficult to change eating patterns. This is important given that mice undergoing the IF 5:2 regimen consumed the same calories as the A/L fed mice. Importantly, the IF 5:2 regimen is also more feasible than the more severe every-other-day fasting (EODF) regimen.

At a molecular level, we demonstrated that PPARα and the GR-PCK1 axis acted in concert as executors of the beneficial effects of the hepatic fasting response. This transcriptional program is activated by NEFAs and GCs, which require a prolonged fasting period comprising of at least 24 hours, as only glucagon-induced glycogenolysis is activated with short-term fasting (Goldstein et al., 2017). In agreement, we also showed that 24 consecutive hours of fasting is more beneficial than 12 hours carried out twice per week against NASH development. Furthermore, a recent study showing mild caloric restriction to improve healthspan in humans found that PPARα-mediated fatty acid oxidation to be one of the most highly upregulated signatures compared to baseline levels, highlighting that many of these beneficial dietary regimens may eventually converge at similar molecular pathways (Spadaro et al., 2022). Notably, we also demonstrate the possibility of mimicking some aspects of the hepatic fasting response, at least at a molecular level, by PPARα agonism with pemafibrate. PPAR agonism is being intensively studied as a possible treatment for NASH. Interestingly, a recent double-blinded, placebo-controlled, randomized multicenter, phase trial demonstrated that pemafibrate significantly decreased liver stiffness (measured by Magnetic Resonance Elastography), serum ALT and liver fibrosis markers as well as lipid parameters compared to placebo controls (Nakajima et al., 2021). This was in agreement with another recent phase 2b, double-blind, randomized, placebo-controlled trial (NATIVE) demonstrating that high-dose (1200 mg) lanifibranor (pan-PPAR agonist) for 24 weeks improved various outcomes of NASH including a reduction in the SAF-A score without worsening of fibrosis as well as a reduction in serum liver damage markers and the majority of lipid, inflammatory and fibrosis biomarkers (Francque et al., 2021). The effectiveness of PPAR agonism in NASH reversal is also supported by multiple pre-clinical mouse studies using either PPARα agonists (*e.g.,* pemafibrate, fenofibrate or Wy-14,643) or the pan-PPAR agonist lanifibranor (Ip et al., 2004; Lefere et al., 2020; Møllerhøj et al., 2022; Sasaki et al., 2020). Notably, our data indicate that it might be possible to further potentiate the effects of PPAR agonism through combined activation of the GR-PCK1 axis, as suggested in our study, but pharmacological agents against PCK1 are not currently available. In line, a recent report highlighted that deletion of *Pck1* exacerbates NASH via activation of a PI3K/AKT/PDGF axis (Ye et al., 2023).

Collectively, given that there is still no pharmacological agent approved for effective NASH treatment, the IF 5:2 regimen may be a cost-effective yet potent intervention for NASH. In accordance, multiple clinical trials have either been conducted or are undergoing to test the role of various IF regimens in NAFLD. In this regard, it was recently shown that a 5:2 intermittent caloric restriction schedule similar to intermittent fasting was highly effective in treating early NAFLD with reduced body weight, liver steatosis and liver stiffness compared to the current standard of care (Holmer et al., 2021). These results are highly promising and further provide evidence to test the role of the IF 5:2 regimen in patients with established NASH. Given the compliance issues with IF however, it is crucial to better understand the molecular mechanisms underpinning the fasting response. In this line, further studies aiming to more faithfully mimic the hepatic fasting response by combined activation of PPARα and PCK1 through the generation of novel PCK1 agonists may yield even greater results.

## Acknowledgements

We are thankful for Corinna Leuchtenberger, Anna Hartley, Katharina Mauel, Axel Szabowski and other members of the M. Heikenwalder’s laboratory for their support, comments and other contributions for this project. We are thankful for Thomas Engleitner and Roland Rad for their support. We are thankful for Max Zimmermann for the aid in kinetic modelling. We are grateful for the Proteomics Core Facility and the Genomics Core Facility at the DKFZ for carrying out the proteomics and the RNA sequencing. We thank Werner Siemens Imaging Center and Werner Siemens Foundation for access to NMR spectroscopy equipment, Miriam Owczorz and Aditi Kulkarni for excellent technical support. Figure schemes were created with BioRender.com.

## Funding

European Research Council (ERC) Consolidator grant (HepatoMetaboPath) (MH) ERC POC (Faith) (MH)

The Helmholtz Future topic Inflammation and Immunology, Zukunftssthema

‘Immunology and Inflammation’ (ZT-0027) (MH) SFB/TR 209 project ID 314905040 (MH, TL) SFB 1479 (Project ID: 441891347) (MH, MR) SFB/TR 179 Project ID 272983813 (MH, SG)

The Rainer Hoenig Stiftung (MH)

Research Foundation Flanders (FWO) under grant 30826052 (EOS Convention MODEL-IDI) (MH)

Seed funding from HI-TRON (MH)

German Research Foundation (DFG) as part of the Excellence Strategy of the federal and state governments — EXC 2180 — 390900677 (NPM)

Bristol Myers Squibb (BMS) company (NS, MMK) DKFZ Clinician Scientist Program (MR)

Dieter Morszeck Foundation (MR) German Cancer Aid grant #70112720 (SR)

## Author contributions

SG, AA, AJR and MH conceived and designed the project. SG and AA conceived, designed, performed and analyzed experiments. JEBA, VJ, PR, XL, EF performed and analyzed experiments. N.S and MMK analyzed proteomics and transcriptomics data. LZ and CT carried out metabolomics and analyzed the metabolomics data. JK, MR, TS, KSK, SP, UR, FM aided with experiments. JH, DH and TM performed liver tissue processing and subsequent immunohistological staining. AJR and NPK provided critical input. TL carried out NAFLD activity score. SH and SR provided human samples. JS analyzed experiments. SG and MH supervised the project. MH secured funding. SG wrote the initial draft of the manuscript with AA and MH providing critical input. All authors provided feedback.

## Competing interests

AJR receives funding for a research project from Boehringer-Ingelheim. All other authors declare that they have no competing interests.

## Data and materials availability

All data are available in the main text or the supplementary materials. Proteomics, metabolomics, and mRNA sequencing data will be deposited in a public database with the published manuscript.

## Supplementary Figure Legends

**Figure S1. Mice under intermittent fasting show resistance against diet-induced obesity and NASH upon Western diet feeding**. 8-week-old male C57BL/6J mice were fed with either normal chow diet or a Western diet (WD) for 32 weeks to induce NASH. The control groups (Chow or WD) were given *ad libitum* (A/L) access to food and water. In contrast, a group of mice fed WD underwent an IF (5:2) regimen involving two non-consecutive days of fasting per week with each fasting cycle lasting 24 hours. **A.** Bodyweight after 32 weeks of diet feeding in mice. Chow A/L (n=5), WD A/L (n=12) and WD IF (5:2) (n=12). **B**. Total fat mass in grams determined by echoMRI at 28 weeks post diet-start. Chow A/L (n=5), WD A/L (n=12) and WD IF (5:2) (n=12). **C and D.** Total lean mass in percentage and grams determined by echoMRI at 28 weeks post diet-feeding. Chow A/L (n=5), WD A/L (n=12) and WD IF (5:2) (n=12). **E and F**. Epididymal white adipose tissue (eWAT) and inguinal white adipose tissue (iWAT) in grams at 32-weeks post diet-feeding, respectively. Chow A/L (n=5), WD A/L (n=12) and WD IF (5:2) (n=12). **G.** Liver weight to body weight ratio in percentage at 32-weeks post diet-start. Chow A/L (n=5), WD A/L (n=12) and WD IF (5:2) (n=12). **H.** Representative immunofluorescence images for CD8, PD1 and DAPI from C57BL/6J mice fed a ND or WD under A/L conditions or WD with IF regimen for 32-weeks. Data are expressed as mean ± SEM. Statistical significance was calculated using one-way analysis of variance with Tukey’s multiple comparison test. ** P < 0.01, *** P < 0.001, **** P < 0.0001. N.S: non-significant. Scale bar, 50µm.

**Figure S2. Fasting-mediated benefits are not calorie-dependent but rather on the length of the fasting cycle. A-J.** Indirect calorimetry was carried out in 8-week-old male C57BL/6J mice to assess food and water intake as well as whole body metabolism. Mice were fed either a normal chow diet or a Western diet (WD). The control groups (Chow or WD) were given *ad libitum* (A/L) access to food and water. Mice under IF (5:2) regimen were fasted for two non-consecutive days per week with each fasting cycle lasting 24 hours. **A.** Cumulative food intake of two days where mice under IF 5:2 underwent fasting. Chow-fed A/L mice (n=4), WD-fed mice (A/L (n=7) and IF 5:2 (n=7). **B.** Average daily water intake assessed over the span of 7 days. **C.** Daily average respiratory exchange ratio (RER) assessed over the span of 7 days. **D.** Average RER during the two days when mice under IF 5:2 underwent fasting cycles. **E.** Average hourly VO_2_ (ml/h) measured over the span of 7 days. **F.** Average hourly activity counts measured over the span of 7 days. **G.** Average daily VO_2_ on the two days when mice under IF 5:2 underwent fasting cycles. **H.** Average daily VO_2_ measured over the span of 7 days. **I.** Average daily activity on the two days when mice under IF 5:2 underwent fasting cycles. **J.** Average total daily activity counts measured over the span of 7 days. Chow-fed A/L mice (n=4), WD-fed mice (A/L (n=7) and IF 5:2 (n=7). **K-T.** 8-week-old male C57BL/6J mice were fed a choline-deficient high-fat diet (CD-HFD) for 32 weeks to induce NASH. CD-HFD A/L mice were given free access to food and water. Mice under the CD-HFD (5:2 - 12h) regimen were fasted for two non-consecutive days per week with each fasting cycle lasting 12 hours from 7pm to 7am. Mice under IF (6:1 - 24h) regimen were fasted for one day per week from 7pm to 7pm with each fasting cycle lasting 24 hours. **K.** Body weight at 32-weeks post diet-start. CD-HFD A/L (n=11), CD-HFD IF (5:2 - 12h) (n=11) and CD-HFD IF (6:1 - 24h) (n=12). **L.** Total fat mass in percentage determined by echoMRI at 28 weeks post diet-start. CD-HFD A/L (n=11), CD-HFD IF (5:2 - 12h) (n=6) and CD-HFD IF (6:1 - 24h) (n=12). **M.** Epididymal white adipose tissue (eWAT) weight in grams at 28 weeks post diet-start. CD-HFD A/L (n=10), CD-HFD IF (5:2 - 12h) (n=11) and CD-HFD IF (6:1 - 24h) (n=12). **N.** Inguinal white adipose tissue (iWAT) weight in grams at 28 weeks post diet-start. CD-HFD A/L (n=9), CD-HFD IF (5:2 - 12h) (n=11) and CD-HFD IF (6:1 - 24h) (n=12). **O.** Serum levels of total cholesterol at 32 weeks post diet-start. CD-HFD A/L (n=11), CD-HFD IF (5:2 - 12h) (n=6) and CD-HFD IF (6:1 - 24h) (n=12). **P**. 16hr fasting serum glucose levels at 28 weeks post diet-start. CD-HFD A/L (n=11), CD-HFD IF (5:2 - 12h) (n=6) and CD-HFD IF (6:1 - 24h) (n=12). **Q and R**. Absolute liver weight and liver weight to body weight ratio in percentage at 32 weeks post diet-start. CD-HFD A/L (n=11), CD-HFD IF (5:2 - 12h) (n=11) and CD-HFD IF (6:1 - 24h) (n=12). **S and T.** Serum levels of alanine transaminase (ALT) and alkaline phosphatase (ALP) at 32 weeks post diet-start. CD-HFD A/L (n=11), CD-HFD IF (5:2 - 12h) (n=11) and CD-HFD IF (6:1 - 24h) (n=12). Data are expressed as mean ± SEM. Statistical significance was calculated using one-way analysis of variance with Tukey’s multiple comparison test. * P < 0.05, ** P < 0.01, *** P < 0.001, **** P < 0.0001. N.S: non-significant.

**Figure S3. IF (5:2 – 12h) regimen does not affect total food intake.** Indirect calorimetry to assess whole body metabolism and total food/water intake was carried out in 8-week-old male C57BL/6J mice fed a choline-deficient high-fat diet (CD-HFD). CD-HFD A/L mice was given free access to food and water. Mice under CD-HFD (5:2 - 12h) regimen were fasted for two non-consecutive days per week with each fasting cycle lasting 12 hours from 7pm to 7am. **A.** Experimental scheme. **B.** Average 12-hourly food intake in mice assessed over the span of 7 days. AL (n=6) and IF (5:2 – 12h) (n=6). **C.** Cumulative food intake assessed over the span of 7 days. AL (n=6) and IF (5:2 – 12h) (n=6). **D.** Average daily water intake assessed over the span of 7 days. AL (n=6) and IF (5:2 – 12h) (n=6). **E and F.** Average hourly respiratory exchange ratio (RER) and VO_2_ assessed over the span of 7 days, respectively. AL (n=6) and IF (5:2 – 12h) (n=6). **G.** Average RER at the times when mice under IF 5:2 – 12h underwent fasting cycles. AL (n=6) and IF (5:2 – 12h) (n=6). **H.** Average daily RER assessed over the span of 7 days. AL (n=6) and IF (5:2 – 12h) (n=6). **I.** Average VO_2_ at the times when mice under IF 5:2 – 12h underwent fasting cycles. AL (n=6) and IF (5:2 – 12h) (n=6). **J.** Average daily VO_2_ assessed over the span of 7 days. AL (n=6) and IF (5:2 – 12h) (n=6). **K.** Average hourly activity counts assessed over the span of 7 days. AL (n=5) and IF (5:2 – 12h) (n=6). **L.** Average activity counts at the times when mice under IF 5:2 – 12h underwent fasting cycles. AL (n=5) and IF (5:2 – 12h) (n=6). **M.** Average daily activity counts assessed over the span of 7 days. AL (n=5) and IF (5:2 – 12h) (n=6). Data are expressed as mean ± SEM. Statistical significance was calculated using one-way analysis of variance with Tukey’s multiple comparison test. * P < 0.05, ** P < 0.01, *** P < 0.001, **** P < 0.0001. N.S: non-significant.

**Figure S4. IF (6:1 – 24h) regimen does not affect total food intake.** Indirect calorimetry to assess whole body metabolism and total food/water intake was carried out in 8-week-old male C57BL/6J mice fed a choline-deficient high-fat diet (CD-HFD). CD-HFD A/L mice was given free access to food and water. Mice under IF (6:1 - 24h) regimen were fasted for one day per week from 7pm to 7pm with each fasting cycle lasting 24 hours. **A.** Experimental scheme **B.** Average daily food intake in mice assessed over the span of 7 days. AL (n=6) and IF (6:1 – 24h) (n=4). **C.** Cumulative food intake for a week. AL (n=6) and IF (6:1 – 24h) (n=4). **D.** Average daily water intake assessed over the span of 7 days. AL (n=6) and IF (6:1 – 24h) (n=4). **E and F.** Average hourly respiratory exchange ratio (RER) and VO_2_ assessed over the span of 7 days, respectively. AL (n=6) and IF (6:1 – 24h) (n=4). **G.** Average RER at the times when mice under IF 5:2 – 12h underwent fasting cycles. AL (n=6) and IF (6:1 – 24h) (n=4). **H.** Average daily RER assessed over the span of 7 days. AL (n=6) and IF (6:1 – 24h) (n=4). **I.** Average VO_2_ at the times when mice under IF 5:2 – 12h underwent fasting cycles. AL (n=6) and IF (6:1 – 24h) (n=4). **J.** Average daily VO_2_ assessed over the span of 7 days. AL (n=6) and IF (6:1 – 24h) (n=4). **K.** Average hourly activity counts assessed over the span of 7 days. AL (n=6) and IF (6:1 – 24h) (n=4). **L.** Average activity counts at the times when mice under IF 5:2 – 12h underwent fasting cycles. AL (n=6) and IF (6:1 – 24h) (n=4). **M.** Average daily activity counts assessed over the span of 7 days. AL (n=6) and IF (6:1 – 24h) (n=4)). Data are expressed as mean ± SEM. Statistical significance was calculated using one-way analysis of variance with Tukey’s multiple comparison test. * P < 0.05, ** P < 0.01, *** P < 0.001, **** P < 0.0001. N.S: non-significant.

**Figure S5. Beneficial effects of fasting in NASH are dependent on the timing and length of the fasting cycle.** 8-week-old male C57BL/6J mice were fed with either normal chow diet or a Western diet (WD) for 32 weeks to induce NASH. The control groups (Chow or WD) were given *ad libitum* (A/L) access to food and water. Mice under WD were also subjected to distinct fasting regimens. Mice under IF (6:1 - 7pm – 7pm) regimen were fasted for one day per week from 7pm to 7pm with each fasting cycle lasting 24 hours. Mice under IF (6:1 - 7am – 7am) regimen were fasted for one day per week from 7am to 7am with each fasting cycle lasting 24 hours. Mice under (5:2 – 7am – 7am) regimen were fasted for two non-consecutive days per week from 7am to 7am with each fasting cycle lasting 24 hours.**A.** Total fat mass in grams determined by echoMRI at 28 weeks post diet-start. Chow-A/L (n=8), WD-A/L (n=11), WD-IF 6:1 (7pm-7pm) (n=12), WD-IF 6:1 (7am-7am) (n=12) and WD-IF 5:2 (n=11). **B and C.** Epididymal white adipose tissue (eWAT) and inguinal white adipose tissue (iWAT) weight in grams at 32-weeks post-diet feeding, respectively. Chow-A/L (n=8), WD-A/L (n=11), WD-IF 6:1 (7pm-7pm) (n=12), WD-IF 6:1 (7am-7am) (n=12) and WD-IF 5:2 (n=11). **D and E.** Total lean mass in percentage and grams determined by echoMRI at 28 weeks post-diet feeding. Chow-A/L (n=8), WD-A/L (n=11), WD-IF 6:1 (7pm-7pm) (n=12), WD-IF 6:1 (7am-7am) (n=12) and WD-IF 5:2 (n=11). **F.** Fasted blood glucose measured at 28-weeks post-diet start. Chow-A/L (n=8), WD-A/L (n=11), WD-IF 6:1 (7pm-7pm) (n=12), WD-IF 6:1 (7am-7am) (n=12) and WD-IF 5:2 (n=11). **G.** Serum alkaline phosphatase (ALP) measured at 32-weeks post-diet start. Chow-A/L (n=8), WD-A/L (n=10), WD-IF 6:1 (7pm-7pm) (n=12), WD-IF 6:1 (7am-7am) (n=11) and WD-IF 5:2 (n=11). **H.** Liver weight to body weight ratio in percentage at 32-weeks post-diet start. Chow-A/L (n=8), WD-A/L (n=11), WD-IF 6:1 (7pm-7pm) (n=12), WD-IF 6:1 (7am-7am) (n=12) and WD-IF 5:2 (n=11). **I.** Summary of the outcome of fasting regimens in the context of two distinct NASH inducing diets in mice. Data are expressed as mean ± SEM. Statistical significance was calculated using one-way analysis of variance with Tukey’s multiple comparison test. *** P < 0.001, **** P < 0.0001. N.S: non-significant.

**Figure S6. Fasting dampens key signaling pathways involved in NASH and HCC.** 8-week-old C57BL/6J mice were fed either a chow diet or a NASH-inducing diet for 5 weeks in the presence or absence of IF 5:2 fasting regimen where mice underwent two fasting cycles per week on two non-consecutive days with each fasting cycle lasting 24 hours. At the end of the 5 weeks, mice fed under A/L conditions were sacrificed in a fed status whilst mice undergoing IF (5:2) regimen were sacrificed either after a 24-hour fast (fasted state) or following 4 hours of refeeding after a 24-hour fast (refed state). **A.** Western blot analysis from liver homogenates harvested from mice sacrificed under different nutritional states such as fed, fasted or refed. **B.** Principal component analysis (PCA) from proteome analysis of livers harvested in different nutritional states **C.** Gene set enrichment analysis (GSEA) for ‘HALLMARK MTORC1 Signaling’ and ‘HALLMARK Glycolysis’ in fasted mice compared to A/L fed controls.

**Figure S7. Hepatocyte-specific glucocorticoid receptor deletion only partially abrogates the hepatic fasting response. A.** Schematics showing activation of glucocorticoid signaling in the hepatocytes during fed or fasted state. **B.** Experimental scheme. 12-week-old GR^fl/fl^ or GR^HEP^ mice were fed a NASH-inducing Western diet (WD) for 5 weeks in the presence or absence of IF 5:2 fasting regimen where mice underwent two fasting cycles per week on two non-consecutive days with each fasting cycle lasting 24 hours. At the end of the 5 weeks, mice fed under A/L conditions were sacrificed in a fed state whereas mice undergoing IF (5:2) regimen were sacrificed after a 24-hour fast (fasted state). **C.** Levels of blood glucose and serum ketone bodies in mice after sacrifice. GR^fl/fl^ (A/L (n=7) and IF (5:2) (n=9), GR^HEP^ (A/L (n=5) and IF (5:2) (n=7). **D-H.** Relative mRNA expression of key metabolic genes in liver homogenates of mice fed a WD. mRNA expression was normalized to *Rps14* housekeeping gene. GR^fl/fl^ (A/L (n=6) and IF (5:2) (n=8), GR^HEP^ (A/L (n=5) and IF (5:2) (n=7). Data are expressed as mean ± SEM. Statistical significance was calculated using one-way analysis of variance with Tukey’s multiple comparison test. * P < 0.05, ** P < 0.01, *** P < 0.001, **** P < 0.0001. N.S: non-significant.

**Figure S8. Depletion of *Pck1* and *Ppara* together abolishes the hepatic fasting response.** 12-week-old C57BL/6J mice were injected intravenously (i.v) with AAV8 vectors carrying either a scrambled shRNA or shRNAs against *Ppara* and/or *Pck1*. Mice were then divided into two groups with one group being fed a Western diet (WD) for 5 weeks under *ad libitum* (A/L) conditions whilst the other group underwent an IF (5:2) regimen. At the end of the 5 weeks, mice fed under A/L conditions were sacrificed in a fed status whilst mice undergoing IF (5:2) regimen were sacrificed after a 24-hour fast (fasted state). **A.** Levels of blood glucose at the end time-point. **B-D.** Relative mRNA expression of key metabolic genes in liver homogenates of mice. mRNA expression was normalized to *Rps14* housekeeping gene. Scramble (A/L (n=6) and IF (5:2) (n=6), shPpara (A/L (n=5) and IF (5:2) (n=5), shPck1 (A/L (n=7) and IF (5:2) (n=6) and combination (A/L (n=5) and IF (5:2) (n=5). **E.** Principal component analysis from RNA-sequencing of liver homogenates from mice. **F.** Gene Set Enrichment Analysis (GSEA) for ‘HALLMARK Inflammatory Response’ and ‘REACTOME Collagen Formation’ of the indicated groups. **G.** Schematics showing that RNA-sequencing was performed from liver homogenates from the indicated mice. **H and I.** Significantly upregulated or downregulated pathways by GSEA under the indicated comparisons. **J and K.** Epididymal white adipose tissue (eWAT) and inguinal white adipose tissue (iWAT) weight in grams at the end time-point. Scramble (A/L (n=6) and IF (5:2) (n=6)), shPpara (A/L (n=5) and IF (5:2) (n=5), shPck1 (A/L (n=7) and IF (5:2) (n=6) and combination (A/L (n=5) and IF (5:2) (n=5). **L and M.** Absolute liver weight and liver to body weight ratio in percentage at the end time-point. Scramble (A/L (n=6) and IF (5:2) (n=6), shPpara (A/L (n=5) and IF (5:2) (n=5), shPck1 (A/L (n=7) and IF (5:2) (n=6) and combination (A/L (n=5) and IF (5:2) (n=5). **N.** Quantification of the number of F4/80^+^ aggregates in livers of mice. **O.** Representative immunohistochemistry images for F4/80 staining in livers of mice. Data are expressed as mean ± SEM. Statistical significance was calculated using one-way analysis of variance with Tukey’s multiple comparison test. * P < 0.05, ** P < 0.01, *** P < 0.001, **** P < 0.0001. N.S: non-significant. Scale bar, 100µm.

**Figure S9. Activation of PPARα signaling by pemafibrate mimics aspects of the hepatic fasting response in mice. A.** Schematic showing the activation of PPARα signaling by non-esterified fatty acids (NEFAs) or pemafibrate. **B.** Experimental scheme. 8-week-old C57BL/6J mice were injected with either vehicle or pemafibrate and sacrificed in a fed state or a fasted (after a 24-hour fasting) state. **C.** Body weight and levels of blood glucose at the end time-point. Fed (n=5), Fasted (n=6) and pemafibrate (n=6). **D-G.** Relative mRNA expression of key metabolic genes in liver homogenates of mice. mRNA expression was normalized to *Rps14* housekeeping gene. Fed (n=5), Fasted (n=6) and pemafibrate (n=6). Data are expressed as mean ± SEM. Statistical significance was calculated using one-way analysis of variance with Tukey’s multiple comparison test. * P < 0.05, ** P < 0.01, *** P < 0.001, **** P < 0.0001. N.S: non-significant.

**Figure S10. Therapeutic fasting ameliorates established NASH in mice.** 8-week-old C57BL/6J mice were fed with a NASH inducing Western diet for 5-months to induce NASH. At this point, mice were evenly distributed to the different treatment arms. Mice were then continued on the diets under *ad libitum* (A/L) conditions or underwent the IF (5:2) regimen for 4 additional months to assess NASH amelioration. **A.** Body weight at 5-months post diet-start and before starting the IF (5:2) regimen. A/L (n=5) and IF (5:2) (n=6). **B and C.** Epididymal white adipose tissue (eWAT) and inguinal white adipose tissue (iWAT) weight in grams at the end time-point. Chow (A/L (n=4), WD (A/L (n=5) and IF (5:2) (n=6). **D.** Serum levels of alkaline phosphatase (ALP) at the end time-point. Chow (A/L (n=4), WD (A/L (n=5) and IF (5:2) (n=6). **E.** Levels of serum cholesterol at the end time-point. Chow (A/L (n=4)), WD (A/L (n=5) and IF (5:2) (n=6). **F.** Liver to body weight ratio in percentage at the end time-point. Chow (A/L (n=4), WD (A/L (n=5) and IF (5:2) (n=6). **G** Experimental scheme. 8-week-old C57BL/6J mice were fed either a chow-diet or a NASH inducing diet such as CD-HFD to induce NASH. A group of mice under CD-HFD were subjected to an IF (5:2) regimen starting from 5-months post diet-start. **H.** Body weight at 5-months post diet-start and before starting the IF (5:2) regimen. CD-HFD (A/L (n=11) and IF (5:2) (n=15). **I.** Body weight change in mice at the end of the experiment compared to body weight at 5-months post diet-start. **J and K.** Epididymal fat and inguinal fat weights in grams at the end time-point. Chow (A/L n=5), CD-HFD (A/L (n=10) and IF (5:2) (n=12). **L and M.** Levels of serum alanine aminotransferase (ALT) and ALP at the end time-point. Chow (A/L n=5), CD-HFD (A/L (n=9) and IF (5:2) (n=12). **N.** Levels of serum cholesterol at the end time-point. Chow (A/L n=5), CD-HFD (A/L (n=9) and IF (5:2) (n=12). **O.** Representative macroscopic liver pictures and immunohistochemistry images for H/E, Sirius red, MHC II and CD8 staining in livers of mice. **P.** Absolute liver weight at the end time-point. Chow (A/L n=5), CD-HFD (A/L (n=9) and IF (5:2) (n=12). **Q-S.** Quantification of the staining in livers of mice. Data are expressed as mean ± SEM. Statistical significance was calculated using one-way analysis of variance with Tukey’s multiple comparison test. * P < 0.05, ** P < 0.01, *** P < 0.001, **** P < 0.0001. N.S: non-significant. Scale bar, 100µm.

**Figure S11. Therapeutic fasting blunts NASH to HCC transition in mice.** 8-week-old male C57BL/6J mice were fed with either a WD for 5 months or choline-deficient high-fat diet (CD-HFD) for 8 months to induce NASH. At this point, mice were evenly distributed to the different treatment arms. Mice were then continued on the diets under A/L conditions or underwent the IF (5:2) regimen to assess NASH-HCC transition at 15 months post-diet start. **A.** Body weight at 5-months post-diet start and before starting the IF (5:2) regimen. WD (A/L (n=23) and IF (5:2) (n=26). **B.** Levels of serum alanine transaminase (ALT) at 5-months post-diet start and before starting the IF (5:2) regimen. WD (A/L (n=25) and IF (5:2) (n=25). **C.** Body weight at 8-months post-diet start and before starting the IF (5:2) regimen. CD-HFD (A/L (n=27) and IF (5:2) (n=23)). **D.** Levels of serum ALT at 8-months post-diet start and before starting the IF (5:2) regimen. CD-HFD (A/L (n=27) and IF (5:2) (n=23). **E.** Monthly body weight measurement in mice after the start of IF 5:2 regimen. Chow (A/L (n=6), WD (A/L (n=11) and IF (5:2) (n=12). **F.** Epididymal white adipose tissue (eWAT) weight in grams. Chow (A/L (n=11), WD (A/L (n=25) and IF (5:2) (n=27). **G**. Monthly body weight measurement in mice after the start of IF 5:2 regimen. Chow (A/L (n=15), CD-HFD (A/L (n=26) and IF (5:2) (n=23). **H.** eWAT weight in grams. Chow (A/L (n=15), CD-HFD (A/L (n=26) and IF (5:2) (n=23). **I.** Inguinal white adipose tissue (iWAT) weight in grams at the end time-point. Chow (A/L (n=11), WD (A/L (n=24) and IF (5:2) (n=25). **J.** Levels of serum cholesterol at the end time-point. Chow (A/L (n=11), WD (A/L (n=24) and IF (5:2) (n=25). **K.** iWAT weight in grams at the end time-point. Chow (A/L (n=15), CD-HFD (A/L (n=26) and IF (5:2) (n=23)). **L.** Levels of serum cholesterol at the end time-point. Chow (A/L (n=15), CD-HFD (A/L (n=26) and IF (5:2) (n=23). **M-T.** Quantification of the staining in livers of mice. **U.** Representative immunohistochemistry images for H/E, MHC II, F4/80 and CD3 staining in livers of mice in chow or western diet fed mice. **V.** Representative immunohistochemistry images for H/E, MHC II, F4/80 and CD3 staining in livers of mice in chow or CD-HFD fed mice. Data are expressed as mean ± SEM. Statistical significance was calculated using one-way analysis of variance with Tukey’s multiple comparison test. * P < 0.05, ** P < 0.01, *** P < 0.001, **** P < 0.0001. N.S: non-significant. Scale bar, 100µm.

**Figure S12. Intermittent fasting does not affect tumor quality in mice fed a Western diet. A.** Representative immunohistochemistry and quantification of images for glutamine synthetase, Ki67, pS6^S240/S244^, pERK, CHOP and p62 staining in livers of mice fed either chow or Western diet with or without IF (5:2) regimen. Data are expressed as mean ± SEM. Statistical significance was calculated using Chi-square test. * P < 0.05, ** P < 0.01, *** P < 0.001, **** P < 0.0001. N.S: non-significant. Scale bar, 5mm (Glutamine synthetase and CHOP) and 100µm (Ki67, pS6^S240/S244^, pERK and p62).

## Methods

### Mice, diets, IF regimens and treatments

All mice used in this study were C57BL/6J. All mice were kept on a 12-hour light/dark cycle and between 21-23° C room temperature under specific pathogen free barrier conditions within individually ventilated cages with *ad libitum* access to food and water unless stated otherwise. Littermate C57BL6/J mice were purchased from either Janvier Labs, France or Charles River, UK. The *GR^fl/fl^* mice in C57BL6/J background were purchased from Jackson Laboratories. The *GR^fl/fl^* mice were then crossed with *Alb-cre* mice to generate hepatocyte-specific GR knockout mice (*GR^HEP^*). The experiments were performed in accordance with German Law and with the approval of the Regierungspräsidium Karlsruhe and Tübingen (G70/18, G139/19, G34/20, R10/20G).

Eight to twelve-week-old male mice were fed a standard chow diet (Provimi Kliba, Switzerland), Western-Diet (WD, Research Diets; D16022301i: 40% kcal fat, 20% kcal fructose 2% cholesterol) or a Choline-Deficient High-Fat Diet (CD-HFD; Research Diets D05010402i: 45% kcal fat, 0% choline) to assess NASH and/or development of subsequent liver cancer.

For prophylactic NASH related experiments, 8-week-old male C57BL/6J mice were fed either a chow diet or NASH inducing diets (WD or CD-HFD) for 32-weeks unless stated otherwise. Control mice were given *ad libitum* access to food during the course of the experiment. In contrast, mice undergoing intermittent fasting (IF) were deprived of food on the fasting periods depending on the IF regimen. A list of IF regimens tested in the context of this study are summarized below:

| Fasting regimen | Beginning time of the fasting cycle | Ending time of the fasting cycle | # of fasting cycles/week | # of fasting hours/week |
| --- | --- | --- | --- | --- |
| IF 5:2 (PM) | 19:00 | 19:00 | 2 | 48 |
| IF 5:2 (AM) | 07:00 | 07:00 | 2 | 48 |
| IF 5:2 – 12h (PM) | 19:00 | 07:00 | 2 | 24 |
| IF 6:1 (PM) | 19:00 | 19:00 | 1 | 24 |
| IF 6:1 (AM) | 07:00 | 07:00 | 1 | 24 |

For the experiments involving therapeutic intervention of NASH, 8-week-old male C57BL/6J mice were fed either a chow diet or NASH inducing diets (WD or CD-HFD) for 5 months to first induce characteristic features of NASH. At this point, mice were subjected to the IF (5:2) regimen with the indicated diets for 4 additional months and mice were analyzed after 9 months of total NASH diet feeding.

For the experiments involving NASH-HCC transition, 8-week-old male C57BL/6J mice were fed either a chow diet or NASH inducing diets for 5 (WD) or 8 (CD-HFD) months to first induce characteristic features of NASH. Thereupon, mice were subjected to the IF (5:2) regimen with the indicated diets until 15 months of total NASH diet feeding.

For the short-term NASH experiments, 8-week-old male C57BL/6J mice were fed a WD for 5 weeks under either *ad libitum* conditions or undergoing the IF (5:2) regimen. After 5 weeks, the *ad libitum* mice were sacrificed in a fed state (fed harvest) whereas the mice undergoing IF (5:2) regimen were sacrificed after a 24 hour fast (fasted harvest) or were fasted for 24 hours followed by 4 hours of refeeding (refed harvest). The same experimental scheme was used for the *GR^fl/fl^* or G*R^fl/fl^* x *Alb-Cre (GR^HEP^)* experiment.

To mimic the effects of IF using pemafibrate, 8-week-old C57BL/6J mice were either given *ad libitum* access to food or fasted for 24 hours prior to sacrifice. In contract, another group of mice fed under *ad libitum* access to food was given pemafibrate (0.1 mg/kg by oral gavage) twice over the span of 24 hours with the last gavage occurring 4 hours prior to being sacrificed.

For the adeno-associated virus vectors serotype 8 (AAV8) transduction experiment, vectors expressing shRNAs against *Pck1* (Cat. No: shAAV-268220) or *Ppara* (Cat. No. shAAV-269120) under the U6 promoter or the scrambled shRNA as a control (Cat. No: 7040) were purchased from the Vector biolabs (Vector biolabs, United States). The vectors were injected via the tail vein (5 × 10^11^ genome copies/mouse) into 12-week old male C57BL/6J mice. The efficiency of knockdown for *Pck1* or *Ppara* in the liver was confirmed by RT-qPCR and transcriptomics. One week after the injections, mice were given NASH inducing WD for 5 weeks. After 5 weeks, the *ad libitum* mice were sacrificed in a fed state (fed harvest) whereas the mice undergoing IF (5:2) regimen were sacrificed after a 24 hour fast (fasted harvest).

### Measurement of blood/serum parameters

Serum was isolated from blood collected in specialized serum isolation gel tubes (Sarstedt, Z/1.1). Serum liver enzymes such as ALT, ALP and total cholesterol were measured with the DRY-CHEM 500i analyzer (Fujifilm, Japan) according to the manufacturer’s instructions. Corticosterone was measured from flash frozen serum with an ELISA kit (Enzo Life Sciences) according to the manufacturer’s instructions. Similarly, serum NEFAs were measured using NEFA-HR (2) assay kit (Fujifilm, Wako Chemicals) according to the manufacturer’s instructions. Levels of blood ketones (ß-Hydroxybutyrate) were measured using a handheld ketone meter (On Call GK Dual, Swiss Point of Care). Blood glucose was measured using a hand-held glucose analyzer (Accu-Chek Aviva, Roche).

### Body-mass composition and metabolic phenotyping

Whole-body composition was determined non-invasively by EchoMRI^TM^ Whole Body Composition Analyzer (Echo Medical Systems Houston, Texas) with a primary accumulation of 1T4. Body weights of mice were measured prior to EchoMRI analysis to calculate percentage of fat and lean mass. For metabolic phenotyping, mice were subjected to indirect calorimetry in PhenoMaster (TSE systems). Briefly, mice were individually housed during the course of the analysis. After a period of 2-3 days of acclimation, parameters such as food intake, water intake, O_2_ consumption, CO_2_ production, respiratory exchange ratio and total activity were measured consecutively for at least a week with five measurement values every hour. Later, analysis of covariance (ANCOVA) was conducted using SigmaPlot (14.0) to rule out the bias that body weight could affect the measured parameters.

### *In vivo* [^18^F]FDG-PET

Dynamic PET acquisitions were performed on a dedicated small animal microPET scanner (Siemens Healthcare, Knoxville, USA) for up to 60 min. after i.v. injection of 12.25±0.25 MBq [^18^F] Fluorodeoxyglucose ([^18^F] FDG). Mice were kept anesthetized using 1.5-2% isoflurane in oxygen and the blood glucose level was assessed at the start of the scan (HemoCue glucose system, HITADO GmbH, Möhnesee, Germany). Afterwards, magnetresonance imaging (MRI) was performed *for anatomical reference using a T2-weighted TurboRARE sequence (BioSpec 70/30, Bruker, Ettlingen, Germany).* PET data were reconstructed applying an OSEM3D algorithm (Inveon Acquisition Workplace, Siemens, Knoxville, TN, USA) with pixel size ∼ 0.38 cm. After coregistration, volumes of interest (VOIs) were placed in the aortic arc, liver, gut, brain and biceps muscle (PMOD version 4.203, Zurich, Switzerland) and tissue activity curves were calculated. Compartmental modeling of the PET data was performed using the method developed by Garbarino et al.

### Immunoblot analysis

Liver homogenates were prepared in 1x RIPA buffer (Cell Signaling Technology), containing protease inhibitor cocktail and phosphatase inhibitor cocktail (ThermoFisher Scientific) according to manufacturer’s recommendations. Equal amounts of protein was denatured at 95°C for 5 min in 1x Laemmli buffer containing 5% β-mercaptoethanol and separated in 4-15% mini-PROTEAN TGX Precast Protein Gels (Biorad) by electrophoresis and blotted onto Nitrocellulose membrane (ThermoFisher Scientific) by wet electroblotting (Mini Trans-Blot, Bio-Rad). The membranes were blocked with 5% BSA or 5% milk in TBS-T for at least 1 hr. at RT. Primary antibodies enlisted in table supplemental #1 were incubated at 4 °C overnight shaking. Incubation with the secondary antibody such as HRP-anti-mouse IgG and HRP-anti-rabbit IgG was performed according to manufacturer’s instruction. The protein detection was carried out using the Clarity Western ECL Substrate (Bio-Rad) in conjunction with the ChemiDoc Touch imaging equipment (Bio-Rad).

**Table Supplemental 1.** (Western blot antibodies)

| Antigen | Source/clone | Dilution | Provider |
| --- | --- | --- | --- |
| p-AKT (S473) | Rabbit, D9E | 1:2000 | Cell Signaling |
| AKT | Rabbit, C67E7 | 1:1000 | Cell Signaling |
| pS6 (S240/244) | Rabbit, D68F8 | 1:10,000 | Cell Signaling |
| RPS6 | Rabbit, 5G10 | 1:2500 | Cell Signaling |
| p-4EBP1(T37/46) | Rabbit, 236B4 | 1:750 | Cell Signaling |
| 4EBP1 | Rabbit, 53H11 | 1:1000 | Cell Signaling |
| p-P65 (S536) | Rabbit, 93H1 | 1:1000 | Cell Signaling |
| P65 | Mouse, L8F6 | 1:1000 | Cell Signaling |
| p-P38 MAPK (T180/182) | Rabbit, 9211 | 1:1000 | Cell Signaling |
| P38 MAPK | Rabbit, 9212 | 1:1000 | Cell Signaling |
| PCK1 | Rabbit, 16754-1-AP | 1:10,000 | Proteintech |
| PDK4 | Rabbit, 12949-1-AP | 1:4000 | Proteintech |
| γ-Tubulin | Mouse, T6557 | 1:5000 | Sigma-Aldrich |
| Vinculin | Mouse, V9131 | 1:5000 | Sigma-Aldrich |

### Isolation of RNA and quantitative real-time PCR

Bulk RNA was extracted from ∼90 mg of snap-frozen liver in 1 mL TRIzol (Invitrogen) and the RNeasyMini Kit (Qiagen). Following quantity and quality control measured by Nanodrop analyzer (Thermo Scientific), 1μg of purified RNA was subsequently transcribed into cDNA using Quantitect Reverse Transcription Kit (Qiagen) according to the manufacturer’s instructions. For mRNA expression analysis, quantitative real-time PCR was performed in duplicates in 384-well plates using Fast Start SYBR Green Master Rox (Roche) on a QuantStudio™ 5 real-time PCR system (Applied Biosystems, Life Technologies). For a list of all used primers for RT-qPCRs, refer to Supplementary Table 2.

**Table Supplemental 2.**
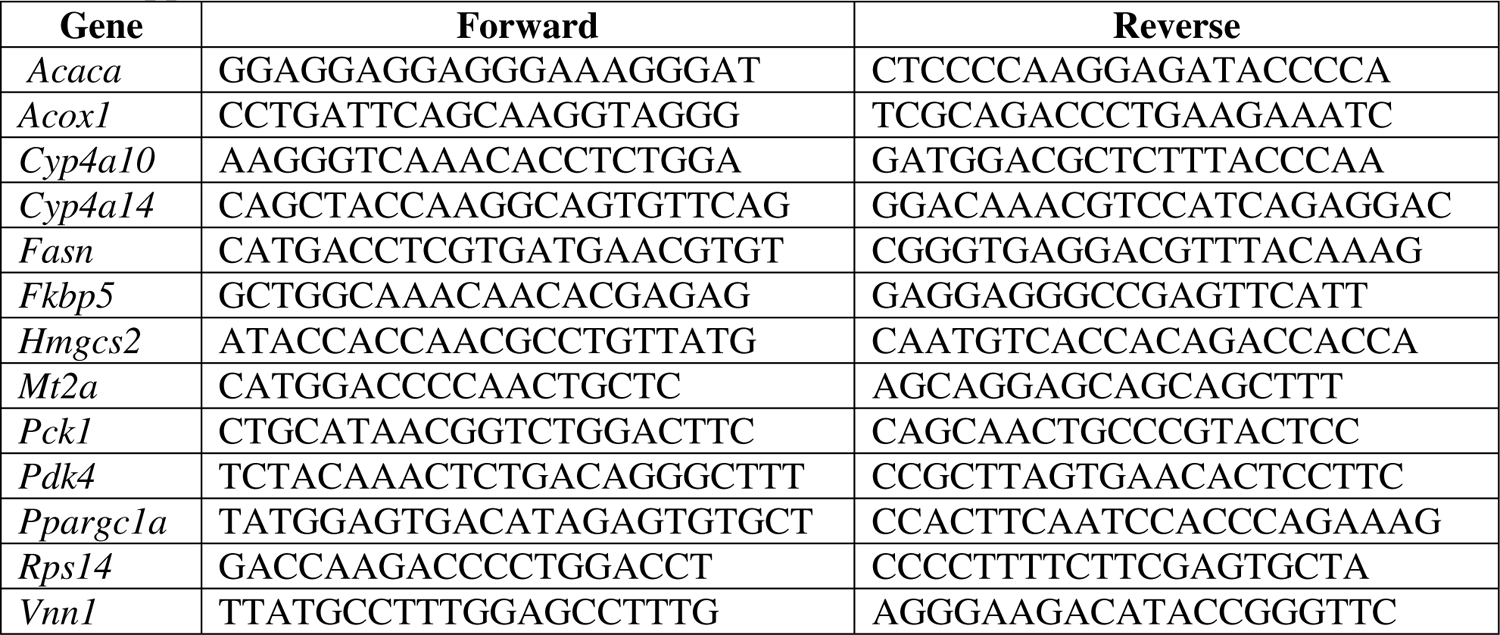
(Primerlist)

### Proteomics - LC-MS/MS sample preparation, measurement and analysis

10 µg of protein sample, isolated as described earlier, was loaded on SDS-gel to run for a short distance of 0.5 cm. After Coomassie staining, the total sample was cut out unfractionated and used for subsequent Trypsin digestion performed on the DigestPro MSi robotic system (INTAVIS Bioanalytical Instruments AG) according to a modified protocol described by Shevchenko et al. digestion (Shevchenko et al., 2006). Digested samples were then loaded on a cartridge trap column, packed with Acclaim PepMap300 C18, 5µm, 300Å wide pore (Thermo Fisher Scientific) and separated in a 180 min gradient from 3% to 40% ACN on a nanoEase MZ Peptide analytical column (300Å, 1.7 µm, 75 µm x 200 mm, Waters) plus an UltiMate 3000 UHPLC system. Data analysis was carried out by MaxQuant (version 1.6.14.0) (Tyanova et al., 2016) using experiment specific databases (Mus musculus; UP000000589; 10090; total entries: 55435) extracted from Uniprot.org under default settings. Identification FDR cut-offs were 0.01 on peptide level and 0.01 on protein level. Match between runs option was enabled to transfer peptide identifications across Raw files based on accurate retention time and m/z. Quantification was done using a label-free quantification approach based on the MaxLFQ algorithm (Cox et al., 2014). A minimum of two quantified peptides per protein was required for protein quantification. A minimum of 2 quantified peptides per protein was required for protein quantification. Data was further processed by in-house compiled R-scripts to plot and filter data.

All differential expression analysis for protein expressions were done using the *test_diff* function in “DEP” (1.16.0) package from Bioconductor. Significant proteins were marked with *add_rejections* where alpha was set to 0.05 and lfc (log fold change) was set to 1. Principal components were calculated using all 2936 proteins with the *prcomp* function. Gene Set Enrichment Analysis (GSEA) was performed using the Broad Institute GSEA desktop application (v4.2.1) based on the ranked adjusted p-values obtained from “DEP” comparisons. Gene sets with less than 15 and more than 500 genes were removed. All analysis was done using R (4.1.2) and R studio (1.4.1106).

### Metabolomics - sample preparation, metabolite extraction and analysis

Fresh-frozen liver tissue (30 – 60 mg) was pulverized (CP02 cryoPREP Automated Dry Pulverizer (220V), with pulverizing impact 2, Covaris LLC, MA, USA) and subjected to a two-phase extraction protocol, previously described in Zizmare, L. et al 2022 (Zizmare et al., 2022). Briefly, liver tissue powder was suspended in 300 µL ultrapure methanol and 1 mL *tert*-butyl methyl ether, shortly vortexed and ultra-sonicated for 5 min for optimal metabolite extraction. Two-phase liquid separation was then achieved by the addition of 250 µL of ultrapure water and centrifugation. Polar and lipid phases were separated manually and solvents were evaporated to dryness. ^1^H-NMR spectroscopy-based metabolomics. The polar dry pellet was re-suspended in 200 µ L deuterated phosphate buffer solution adjusted to pH=7.4, containing internal standard, 1 mM 3-(trimethylsilyl) propionic-2,2,3,3-d_4_ acid sodium salt (TSP), vortexed and centrifuged for optimal suspension and removal of undissolved residue. 190 µ L of clear supernatant were then filled in a 3 mm NMR tube. NMR spectra were recorded by 14.1 Tesla ultra-shielded NMR spectrometer (600 MHz proton frequency, Avance III HD, Bruker BioSpin, Ettlingen, Germany) with 5mm probe head 3 mm experimental operating mode setup at 298 K with Carr-Purcell-Meiboom-Gill (CPMG, number of scans = 1024) pulse program. Spectra were processed by Bruker TopSpin 3.6.1. software (Bruker BioSpin, Ettlingen, Germany). Metabolite identification assignments and quantification was performed with ChenomX NMR suite 8.5 (ChenomX Inc, Edmonton, Canada) software, including human metabolome database (HMDB) reference NMR spectra information, and a quantified metabolite concentration table was exported.

Statistical data analysis was performed on the R-based MetaboAnalyst 5.0 platform (https://www.metaboanalyst.ca) (Pang et al., 2021). Metabolite raw concentrations were normalized by the probabilistic quotient normalization (PQN) method to account for dilution effects. Analysis provided principal component analysis (PCA); metabolite heatmap illustrating auto-scaled concentration values and metabolite concentration pattern connectivity visualization, based on unsupervised hierarchical cluster analysis with Euclidean distance measure and Ward clustering method; and quantitative pathway enrichment analysis, based on the small molecule pathway database (SMPDB) 99 metabolite sets, illustrating enrichment ratio (observed hits / expected hits per pathway) and pathway alteration significance based on individual metabolite p-values.

### RNA-sequencing and analysis

Sequencing libraries were prepared using the Illumina TruSeq mRNA stranded Kit following the manufacturer’s instructions by the DKFZ Genomics Core Facility. Briefly, mRNA was purified from 500ng of total RNA using oligo(dT) beads. Then poly(A)+ RNA was fragmented to 150 bp and converted to cDNA. The cDNA fragments were then end-repaired, adenylated on the 3′ end, adapter ligated and amplified with 15 cycles of PCR. The final libraries were validated using Qubit (Invitrogen) and Tapetstation (Agilent Technologies). 2x 100 bp paired-end sequencing was performed on the Illumina NovaSeq 6000 according to the manufacturer’s protocol. At least 58 Mio. reads per sample were generated.

Raw reads were analyzed using *nf-core rnaseq* (3.8.1) pipeline (Nextflow version 22.04.0) with *STAR-RSEM* as the choice for aligner. Reads were mapped to the GRCm38 genome using STAR. RNA-Seq differential expression analysis was performed with “DESeq2” package from Bioconductor as part of this pipeline. GSEA analysis performed with the same parameters as proteomics above. All analysis was done using R (4.1.2) and R studio (1.4.1106).

### Immunohistochemistry and immunofluorescence

Tissues were fixed in paraformaldehyde for 48-hours, thereafter dehydrated and embedded in paraffin. Antigens were detected in 2-µm tissue slices whereas Sirius-Red staining was performed on 7-µm slices as explained before (Gallage et al., 2022). In brief, cuts were deparaffinazed and rehydrated, antigens were retrieved with Bond citrate solution (AR9961, Leica), Bond EDTA solution (AR9640, Leica) or Bond proteolytic enzyme kit (AR9551, Leica). Thereafter, the slides were incubated with primary antibodies (Table 3) in Bond primary antibody diluent (AR9352, Leica) followed by secondary antibodies (Leica) and stained with the Bond Polymer Refine Detection Kit (DS9800, Leica). The slides were scanned with a whole-slide scanner (Aperio AT2, Leica) at 20x. The slides were annotated and analyzed with Aperio ImageScope (ver. 12.4.0.5043, Leica) and Fiji (ImageJ ver. 1.52e, National Institutes of Health USA). The NAFLD activity score (NAS) was carried out by a certified pathologist (Prof. Thomas Longerich).

Immunofluorescence stainigs were done with established primary antibodies (Table 3) followed by the AKOYA Biosciences Opal Fluorophore kit (Opal 520, FP1487001KT and Opal 620, FP1495001KT) and counterstained with AKOYA 10x Spectral DAPI (FP1490) following the manufacturer’s instructions. The slides where imaged with a NanoZoomer S60 Scanner (Hamamatsu) at 20x and analyzed with the NDP.view software (ver. 2.7.25, Hamamatsu).

**Table Supplemental 3.** (IHC and Immunofluorescence antibodies)

| Antigen | Source/clone | Dilution | Provider |
| --- | --- | --- | --- |
| F4/80 | Rat, BM8 | 1:250 | Linaris |
| MHCII | Rat, M5/114.15.2 | 1:500 | Novus Biologicals |
| CD42b | Rabbit, SP219 | 1:200 | Abcam |
| CD8 | Rat, 4SM15 | 1:200 | Invitrogen |
| PD1 | Goat, AF1021 | 1:100 | Biotechne |
| CD3 | Rabbit, SP7 | 1:500 | Invitrogen |
| Glutamine synthetase | Rabbit, ab16802 | 1:1000 | Abcam |
| Ki67 | Rabbit, SP6 | 1:100 | Fisher Scientific |
| pS6-S240/S244 | Rabbit, D68F8 | 1:2000 | Cell Signalling |
| pERK1/2 | Rabbit, D13.14.4E | 1:400 | Cell Signalling |
| CHOP | Rabbit, D46F1 | 1:100 | Cell Signaling |
| p62 | Rabbit, MBL-PM045 | 1:500 | Biozol diagnostica |

### Statistical analyses

Data are expressed as mean ± standard error of mean (SEM) unless otherwise stated. Statistical analysis was performed in Graph Pad Prism 9. Statistical significance for comparison of only two groups was done by Student’s t-test. Statistical significance for comparison of three or more groups with only one variable was calculated using one-way analysis of variance with Tukey’s multiple comparison test. Statistical significance for comparison of four or more groups with two variables (*e.g.,* genotype and dietary regimen) was calculated using two-way analysis of variance with Tukey’s multiple comparison test. Statistical significance for the tumor incidence was carried out using the Fischer’s exact test. A p­value of < 0.05 was considered statistically significant. * P <0.05, ** P< 0.01, *** P<0.001 and **** P<0.0001.

