## Supplementary Figures for "Beneficial effects of intermittent fasting in NASH and subsequent HCC development are executed by concerted PPARα and PCK1 action in hepatocytes"

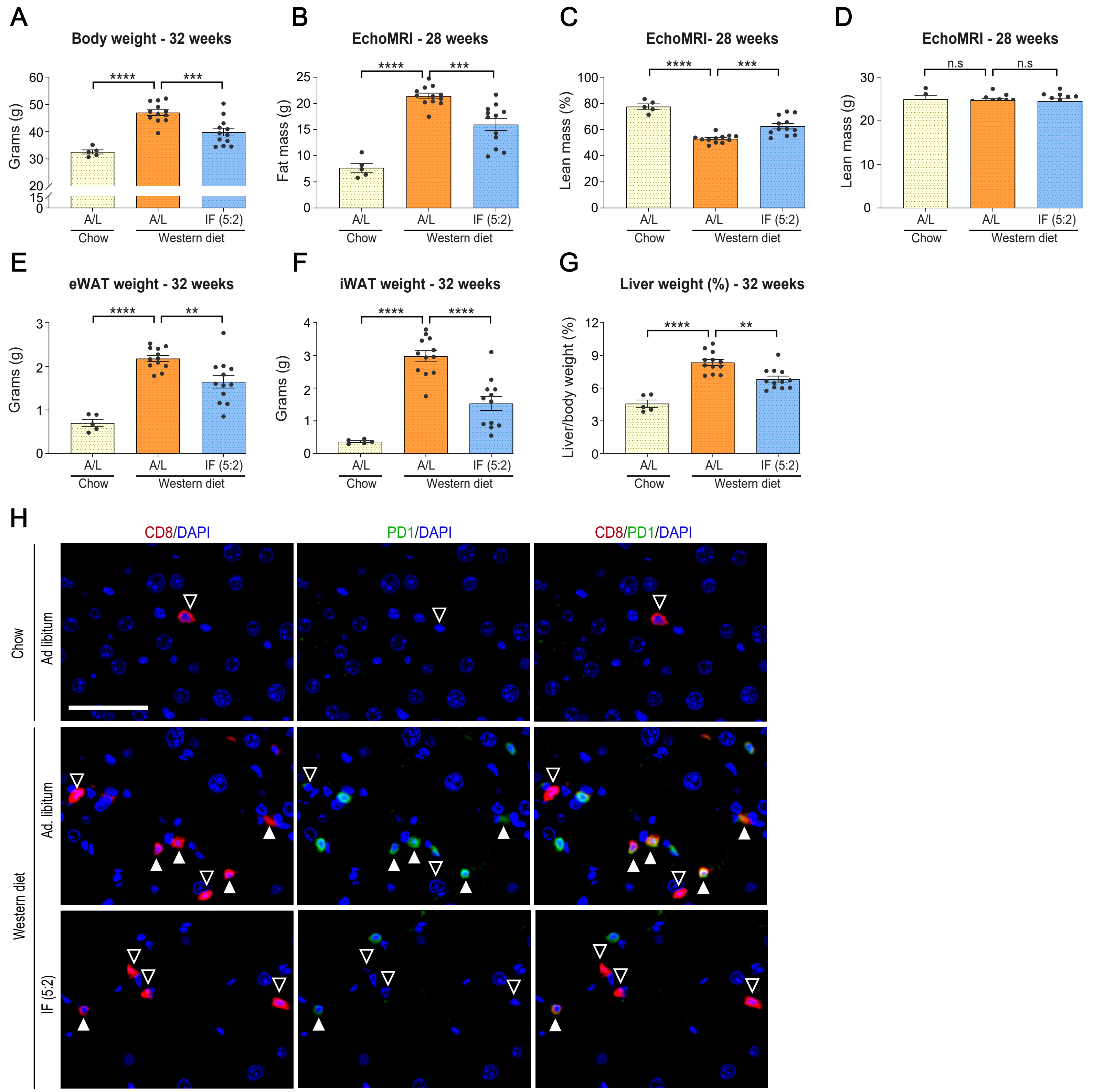


Supplementary Figure. 1

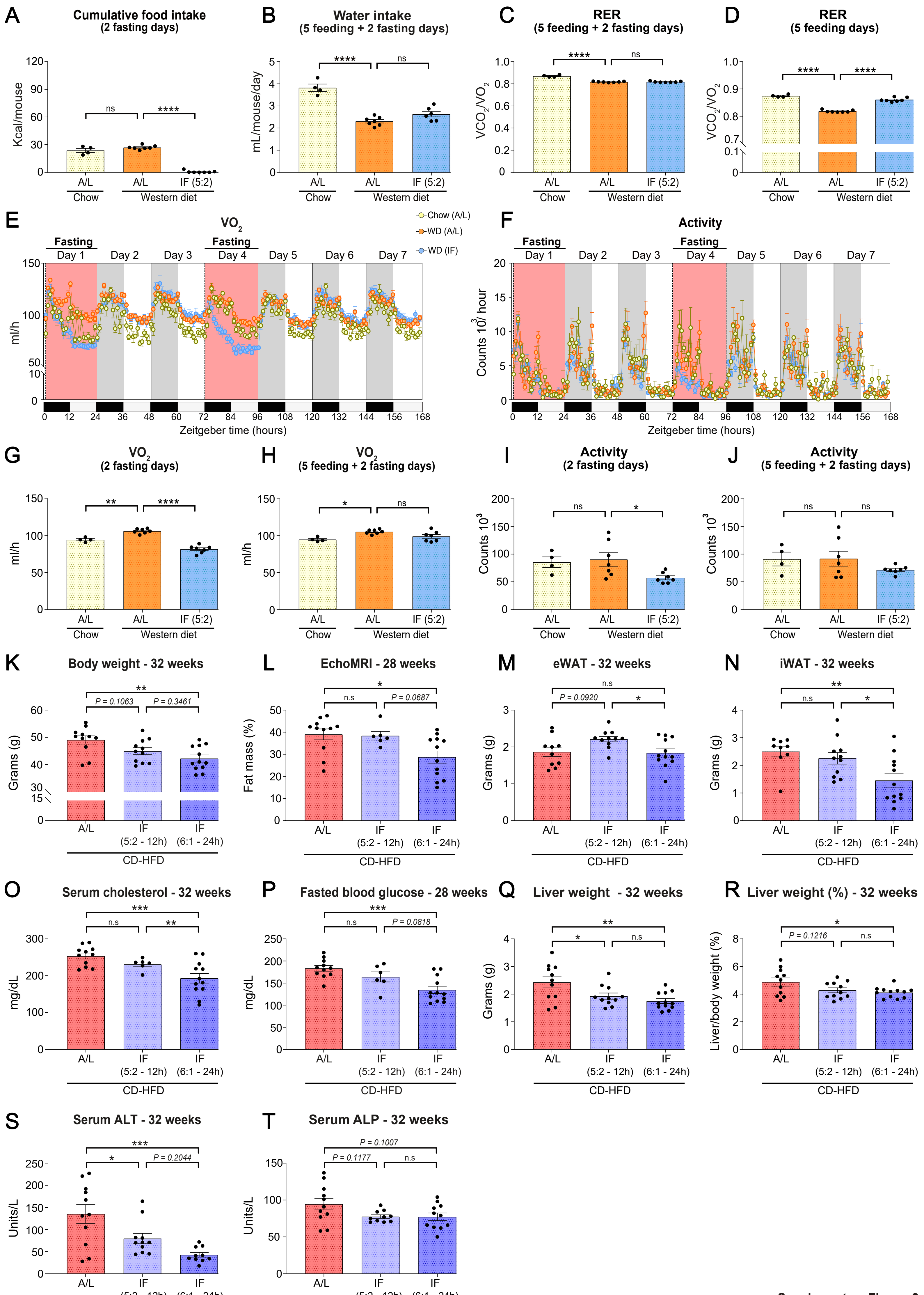


Supplementary Figure. 2

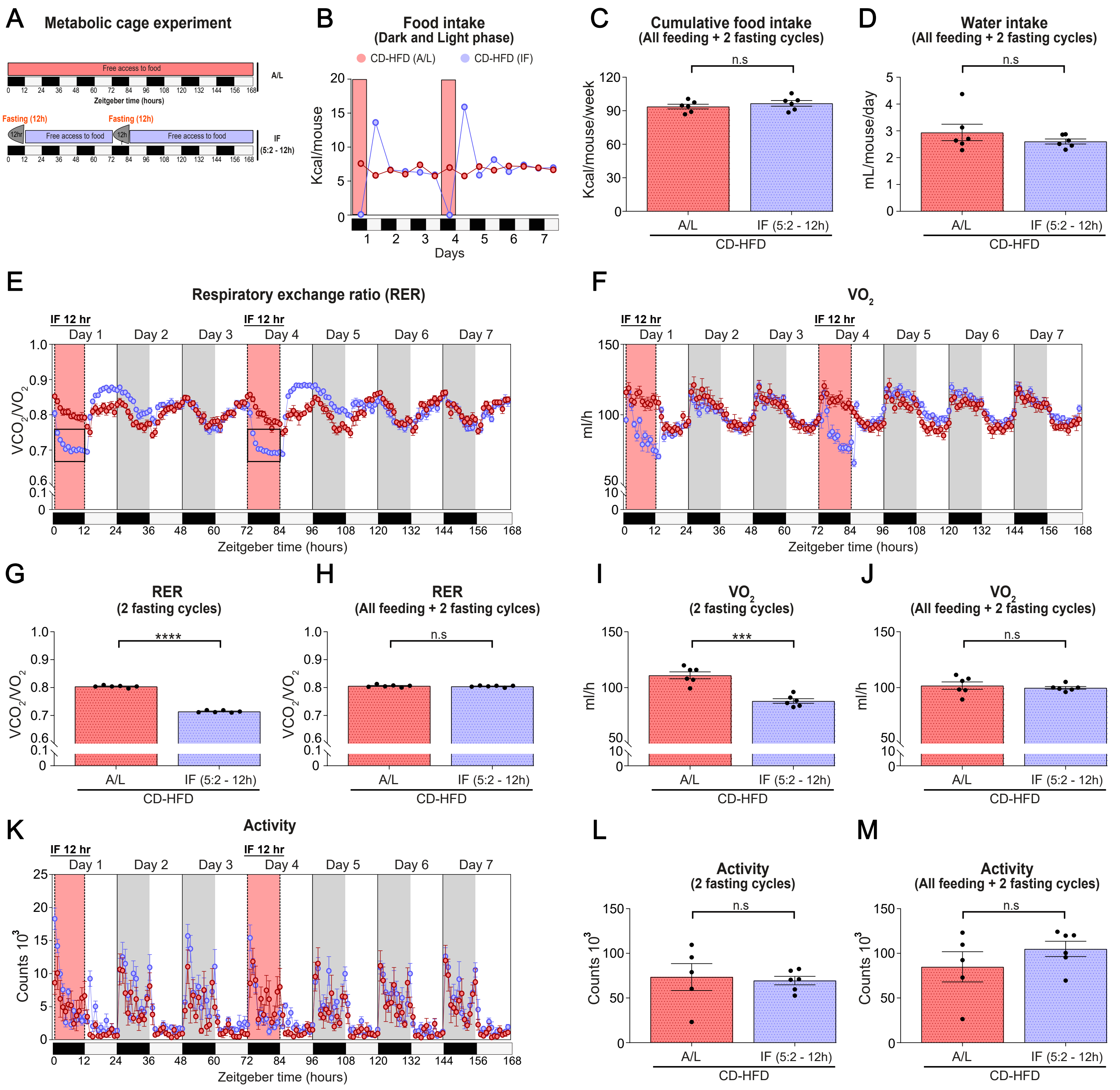


Supplementary Figure. 3

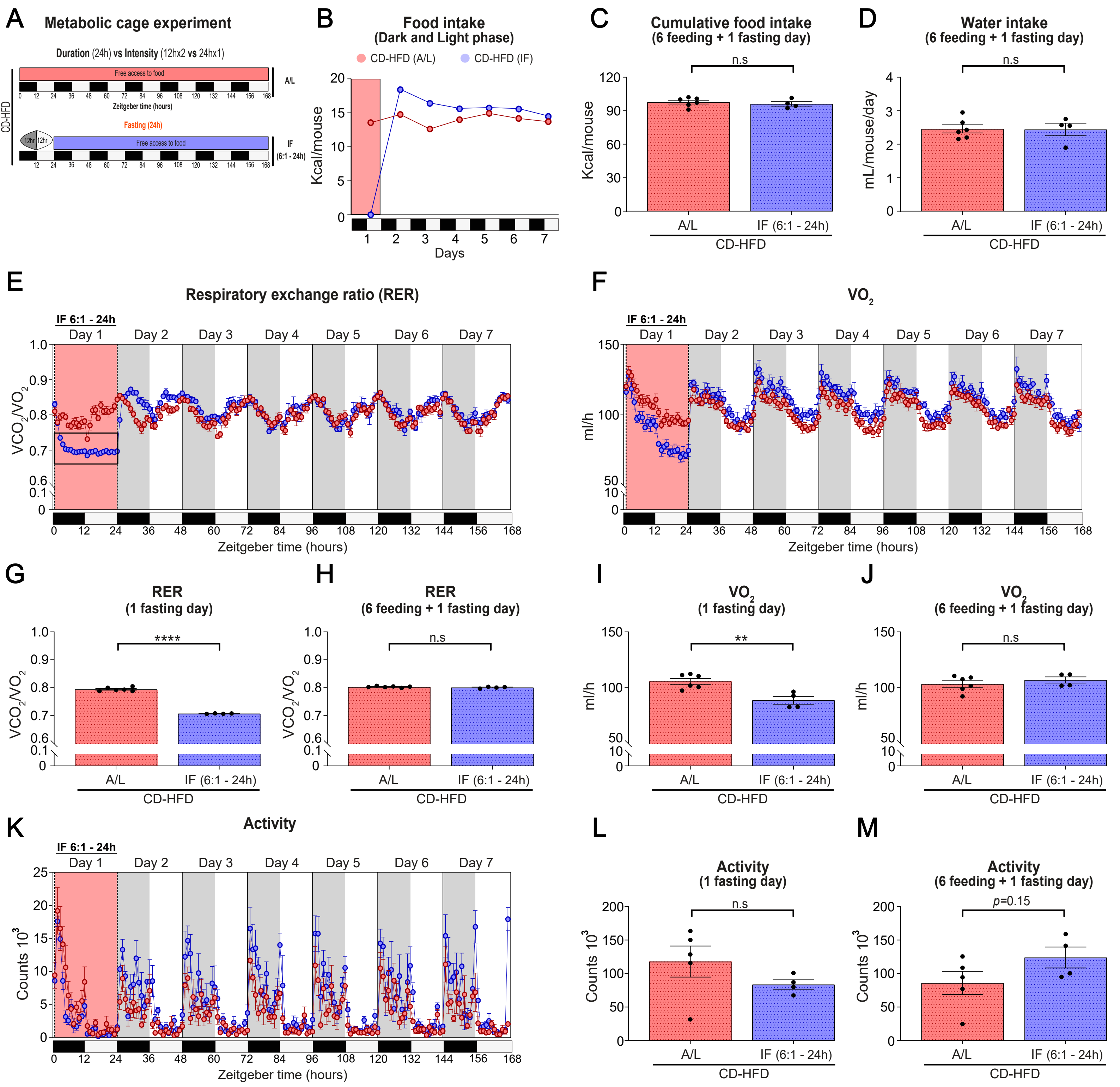


Supplementary Figure. 4

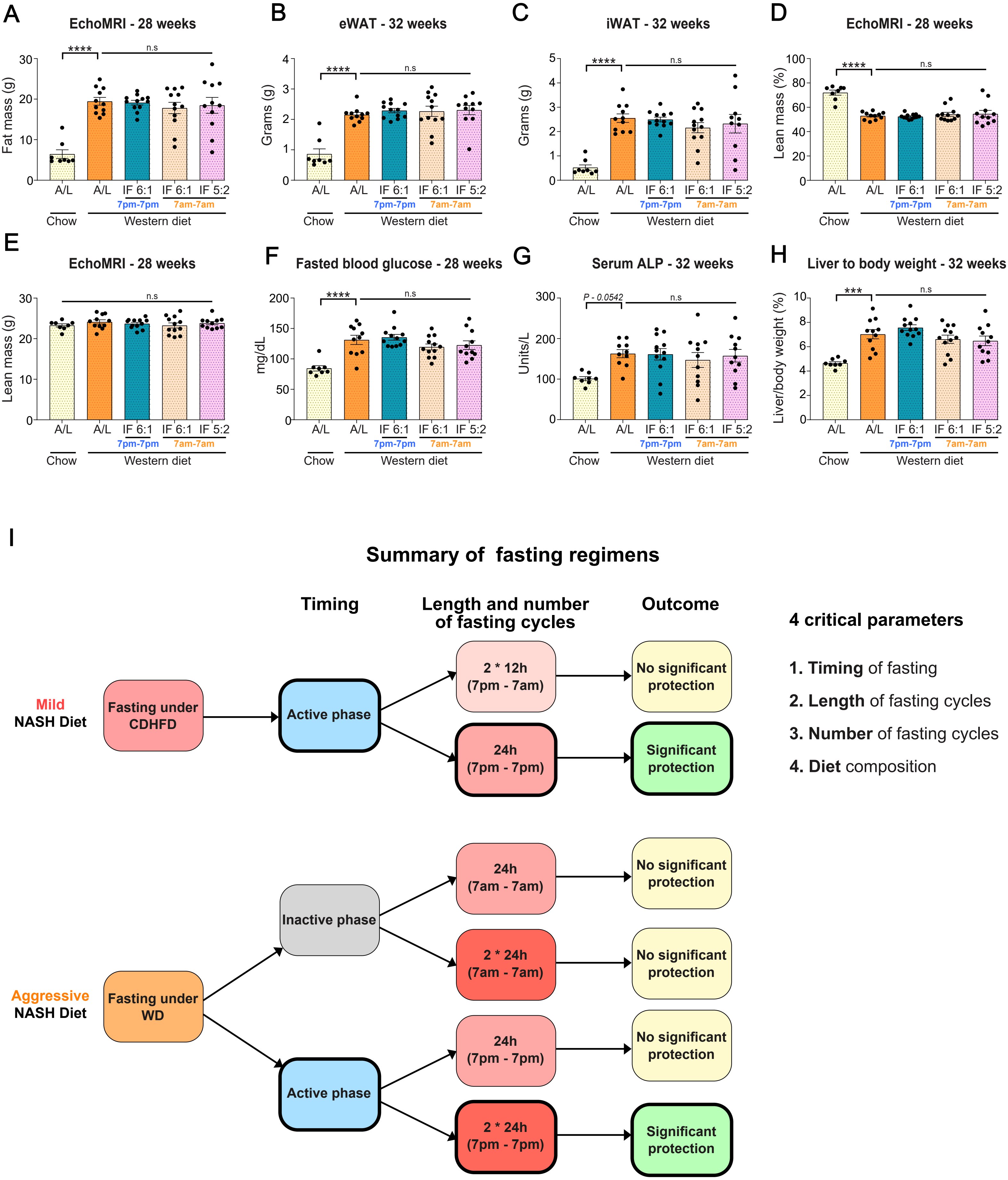


Supplementary Figure. 5

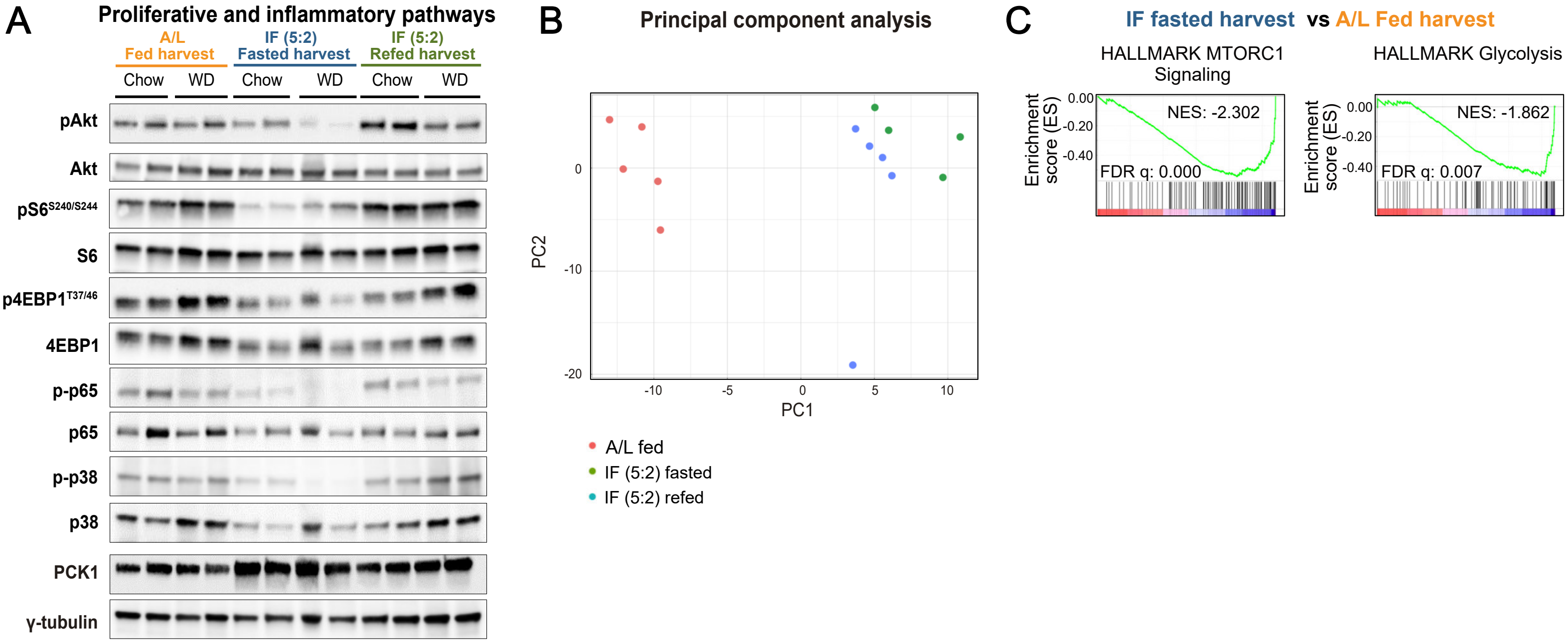


Supplementary Figure. 6

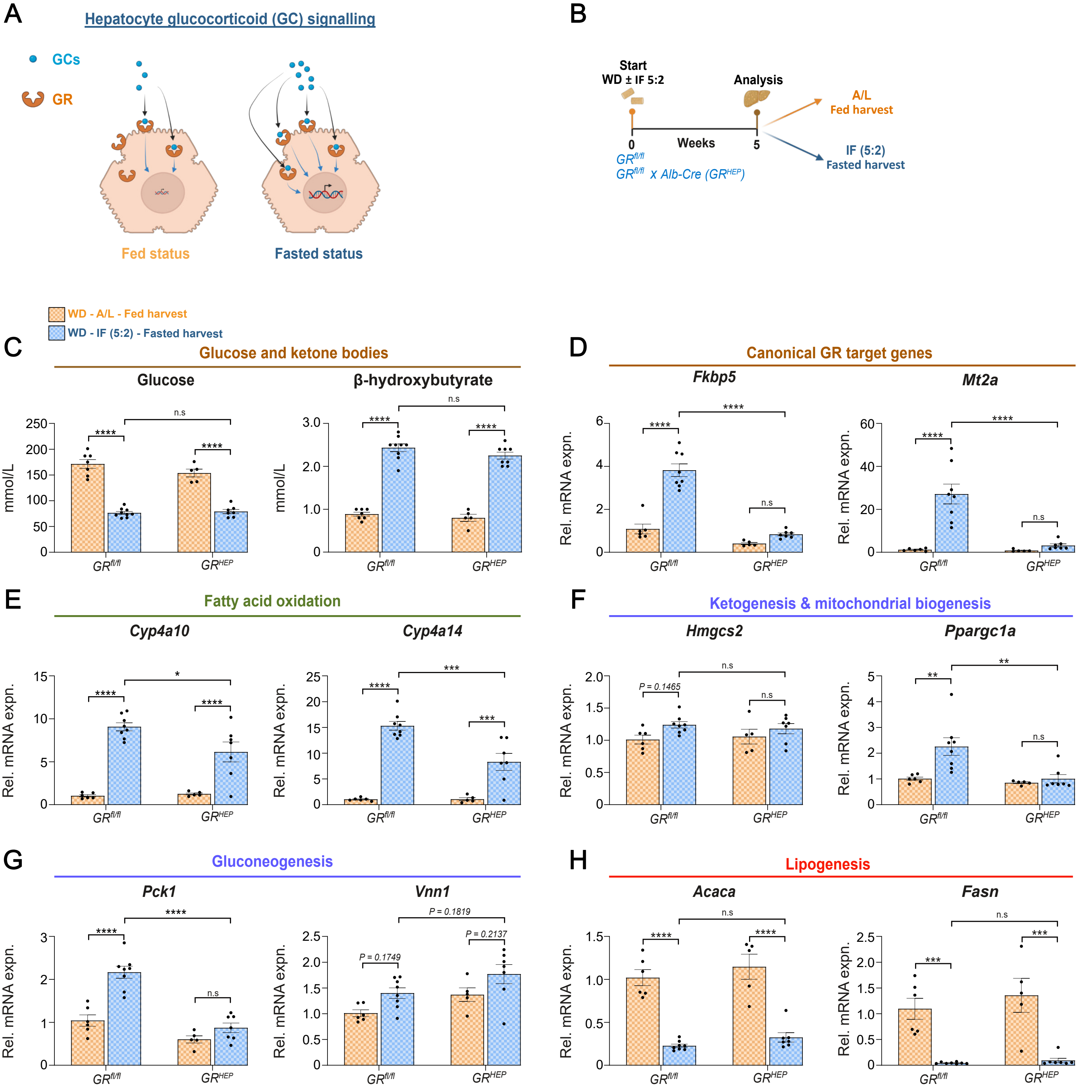
A
Hepatocyte glucocorticoid (GC) signalling
GCs
GR
Fed status
Fasted status
B
Start
WD ± IF 5:2
Analysis
A/L
Fed harvest
IF (5:2)
Fasted harvest
0
Weeks
5
GRfl/fl
GRfl/fl x Alb-Cre (GRHEP)
WD - A/L - Fed harvest
WD - IF (5:2) - Fasted harvest
C
Glucose and ketone bodies
Glucose
β-hydroxybutyrate
mmol/L
D
Canonical GR target genes
Fkbp5
Mt2a
Rel. mRNA expn.
E
Fatty acid oxidation
Cyp4a10
Cyp4a14
F
Ketogenesis & mitochondrial biogenesis
Hmgcs2
Ppargc1a
P = 0.1465
G
Gluconeogenesis
Pck1
Vnn1
P = 0.1749
P = 0.1819
P = 0.2137
H
Lipogenesis
Acaca
Fasn
GRfl/fl
GRHEP
n.s

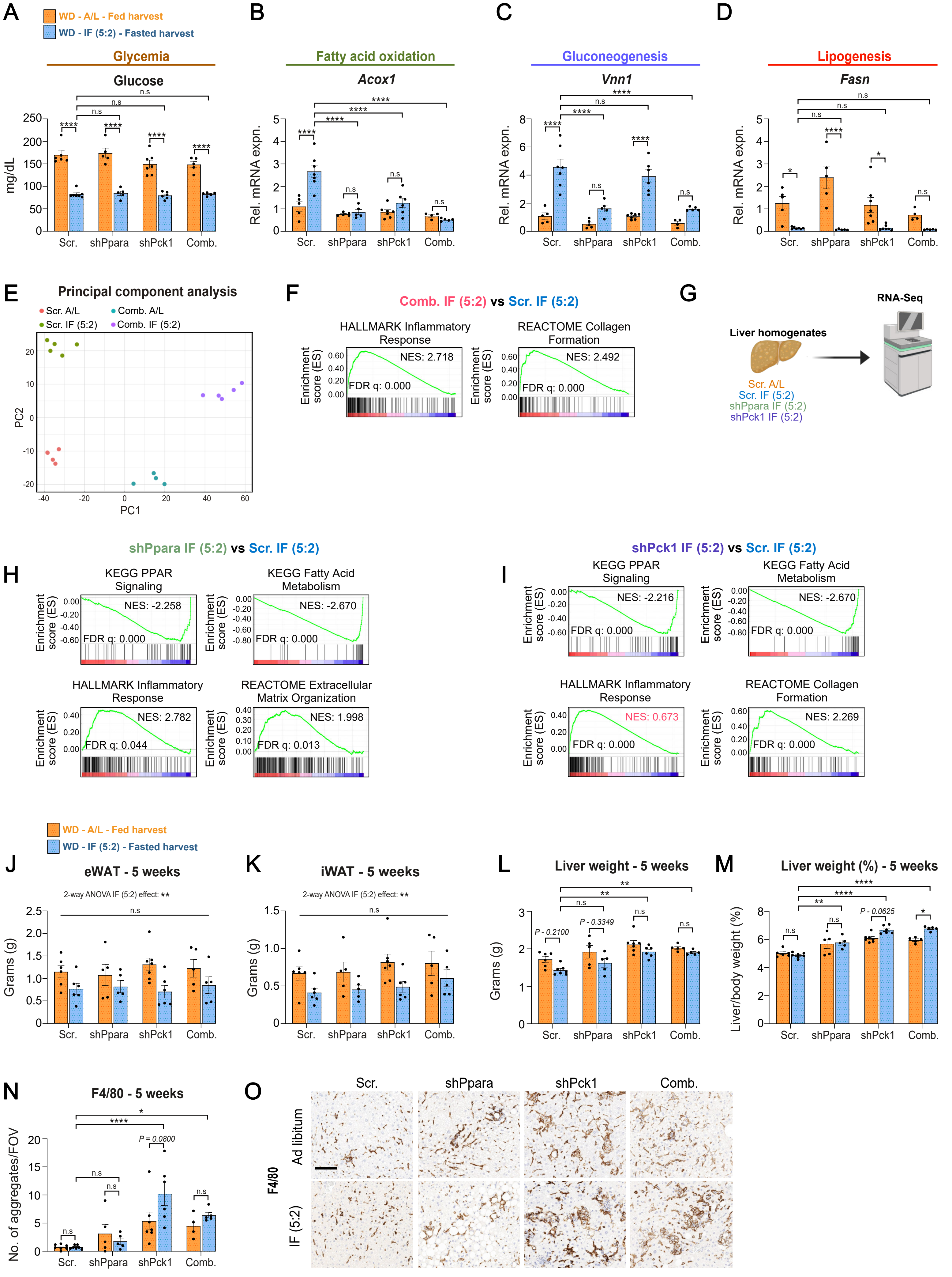

A
WD - A/L - Fed harvest
WD - IF (5:2) - Fasted harvest
Glycemia
Glucose
mg/dL
Scr. shPpara shPck1 Comb.
B
Fatty acid oxidation
Acox1
Rel. mRNA expn.
C
Gluconeogenesis
Vnn1
D
Lipogenesis
Fasn
E
Principal component analysis
Scr. A/L
Comb. A/L
Scr. IF (5:2)
Comb. IF (5:2)
PC1
PC2
F
Comb. IF (5:2) vs Scr. IF (5:2)
HALLMARK Inflammatory Response
NES: 2.718
FDR q: 0.000
REACTOME Collagen Formation
NES: 2.492
FDR q: 0.000
Enrichment score (ES)
G
Liver homogenates
RNA-Seq
Scr. A/L
Scr. IF (5:2)
shPpara IF (5:2)
shPck1 IF (5:2)
shPpara IF (5:2) vs Scr. IF (5:2)
H
KEGG PPAR Signaling
NES: -2.258
FDR q: 0.000
KEGG Fatty Acid Metabolism
NES: -2.670
FDR q: 0.000
HALLMARK Inflammatory Response
NES: 2.782
FDR q: 0.044
REACTOME Extracellular Matrix Organization
NES: 1.998
FDR q: 0.013
shPck1 IF (5:2) vs Scr. IF (5:2)
I
KEGG PPAR Signaling
NES: -2.216
FDR q: 0.000
KEGG Fatty Acid Metabolism
NES: -2.670
FDR q: 0.000
HALLMARK Inflammatory Response
NES: 0.673
FDR q: 0.000
REACTOME Collagen Formation
NES: 2.269
FDR q: 0.000
WD - A/L - Fed harvest
WD - IF (5:2) - Fasted harvest
J
eWAT - 5 weeks
2-way ANOVA IF (5:2) effect: **
Grams (g)
K
iWAT - 5 weeks
2-way ANOVA IF (5:2) effect: **
L
Liver weight - 5 weeks
P - 0.2100
P - 0.3349
M
Liver weight (%) - 5 weeks
Liver/body weight (%)
P - 0.0625
N
F4/80 - 5 weeks
No. of aggregates/FOV
P = 0.0800
O
Scr. shPpara shPck1 Comb.
Ad libitum
IF (5:2)
F4/80

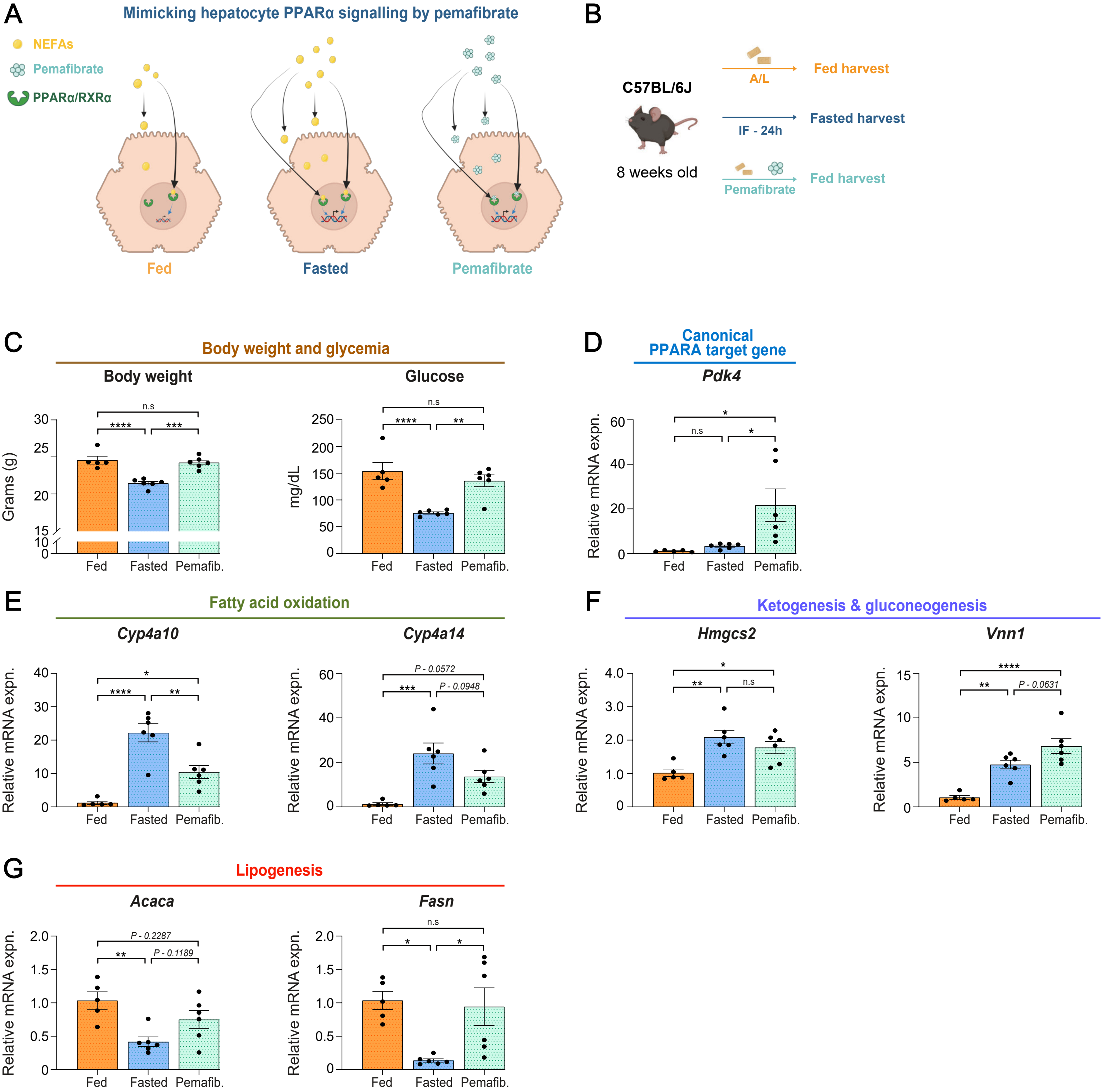
A
Mimicking hepatocyte PPARα signalling by pemafibrate
NEFAs
Pemafibrate
PPARα/RXRα
Fed
Fasted
Pemafibrate
B
C57BL/6J
8 weeks old
A/L
Fed harvest
IF - 24h
Fasted harvest
Pemafibrate
Fed harvest
C
Body weight and glycemia
Body weight
Grams (g)
Glucose
mg/dL
D
Canonical
PPARA target gene
Pdk4
Relative mRNA expn.
E
Fatty acid oxidation
Cyp4a10
Cyp4a14
P - 0.0572
P - 0.0948
F
Ketogenesis & gluconeogenesis
Hmgcs2
Vnn1
P - 0.0631
G
Lipogenesis
Acaca
P - 0.2287
P - 0.1189
Fasn
Fed
Fasted
Pemafib.

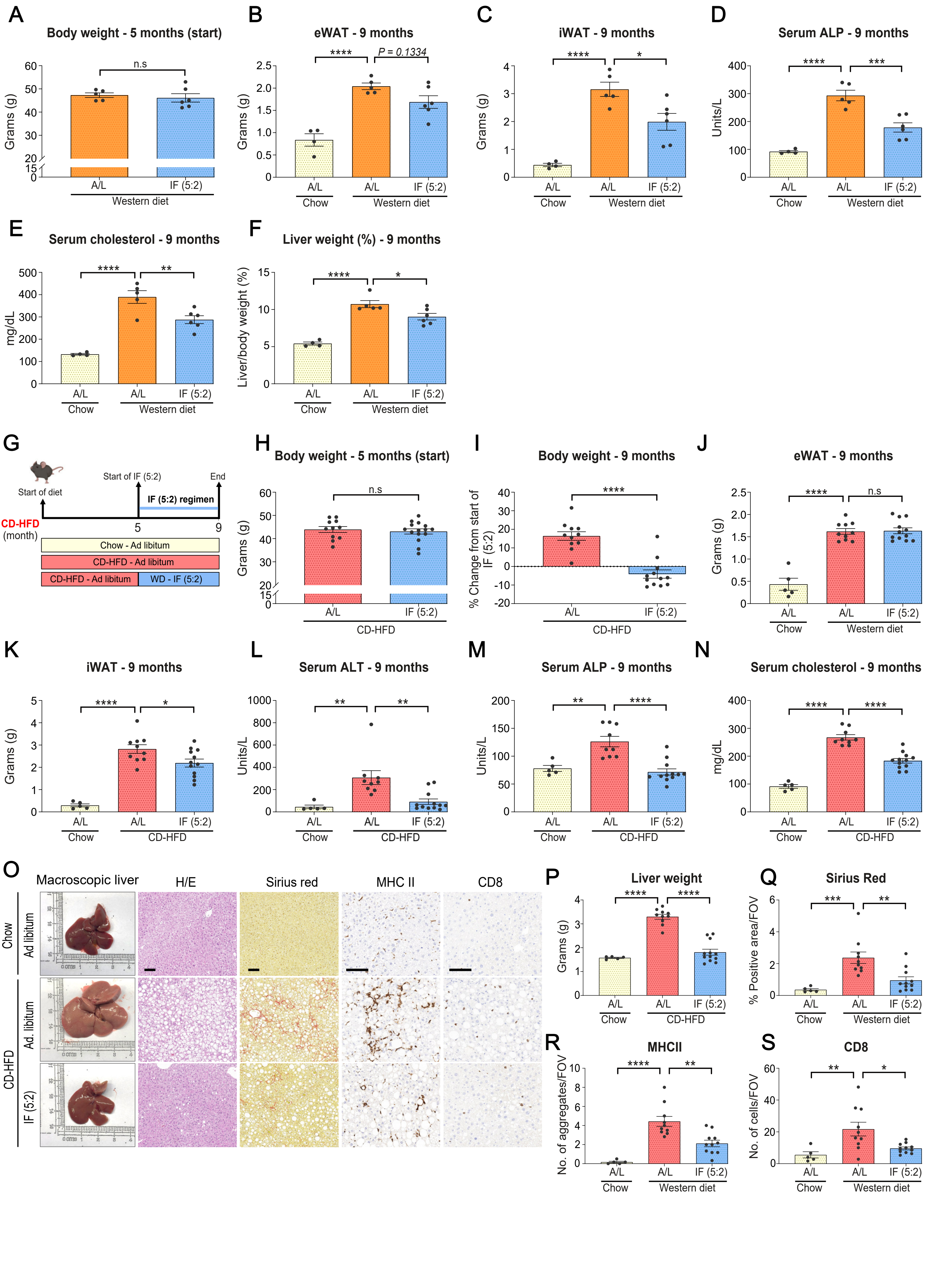


Supplementary Figure. 10

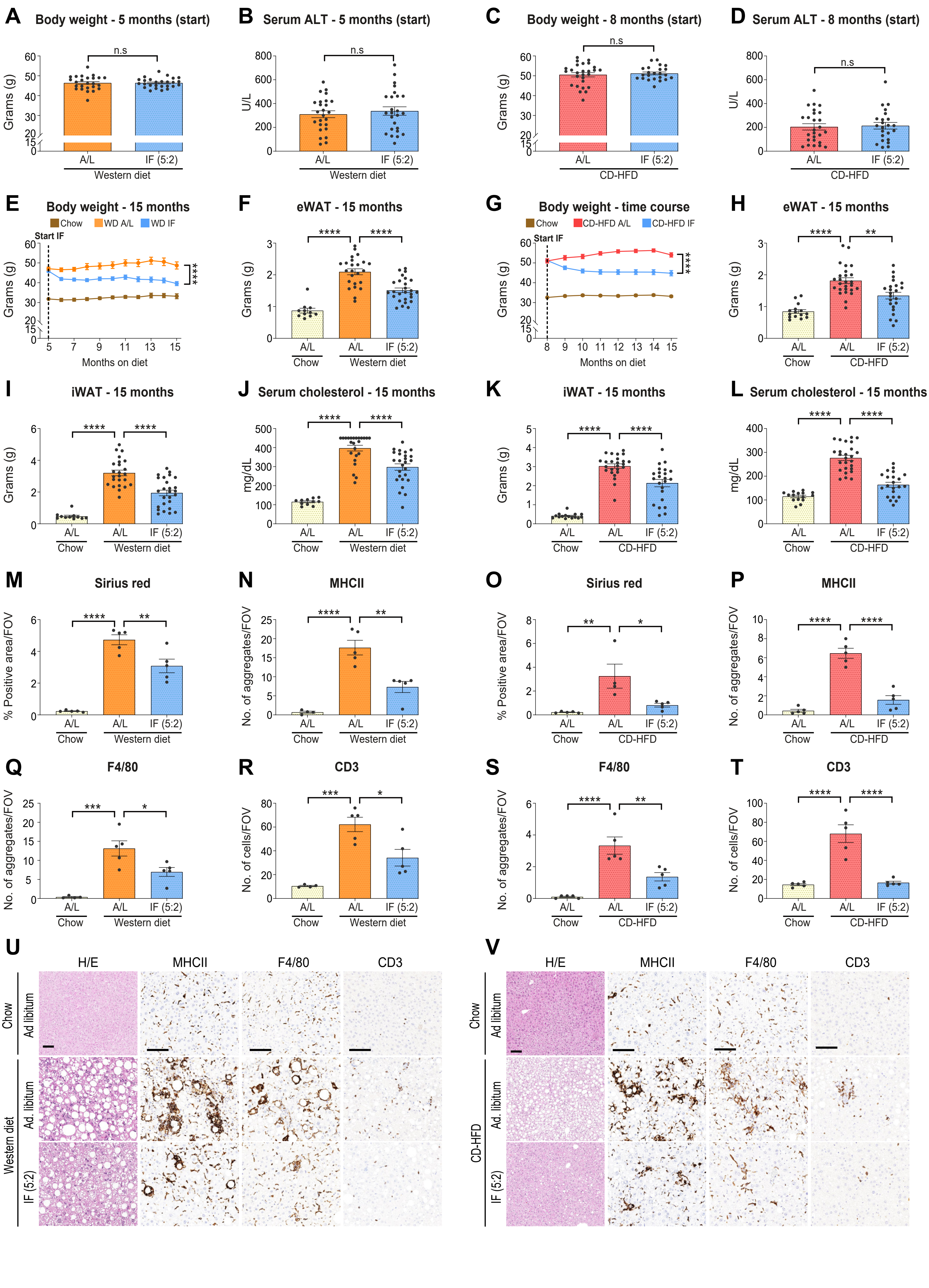


Supplementary Figure. 11

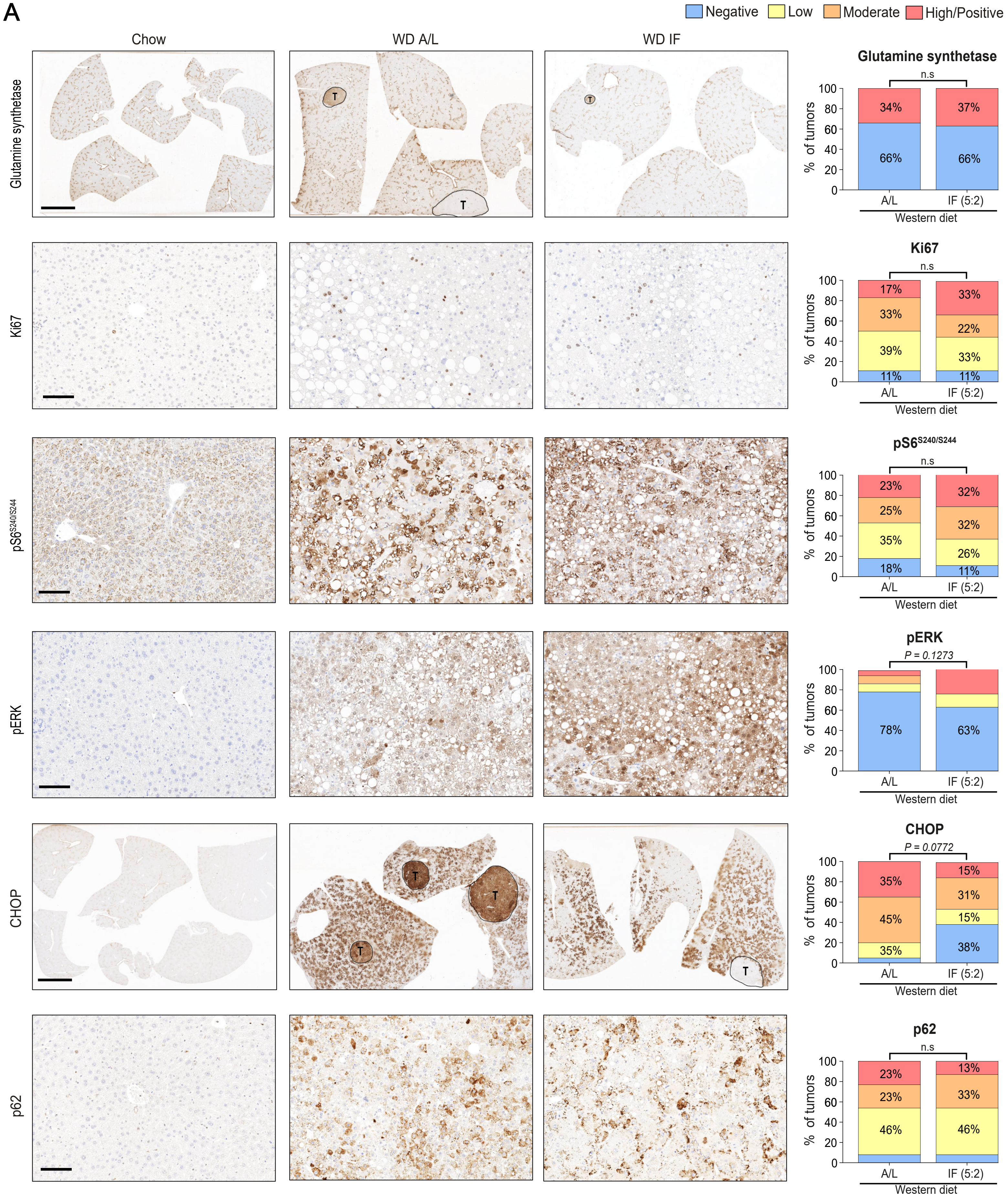
A
Negative
Low
Moderate
High/Positive
Chow
WD A/L
WD IF
Glutamine synthetase
Ki67
pS6 S240/S244
pERK
CHOP
p62
T
Glutamine synthetase
n.s
34%
37%
66%
66%
Ki67
n.s
17%
33%
33%
22%
39%
33%
11%
11%
pS6 S240/S244
n.s
23%
32%
25%
32%
35%
26%
18%
11%
pERK
P = 0.1273
78%
63%
CHOP
P = 0.0772
35%
15%
31%
45%
15%
35%
38%
p62
n.s
23%
13%
23%
33%
46%
46%
% of tumors
A/L
IF (5:2)
Western diet
